# Multi-Omics Analysis of Magnetically Levitated Plasma Biomolecules

**DOI:** 10.1101/2022.09.06.506852

**Authors:** Ali Akbar Ashkarran, Hassan Gharibi, Dalia Abou Zeki, Irina Radu, Farnaz Khalighinejad, Kiandokht Keyhanian, Christoffer K. Abrahamsson, Carolina Ionete, Amir Ata Saei, Morteza Mahmoudi

**Affiliations:** Department of Radiology and Precision Health Program, Michigan State University, East Lansing, MI, USA; Division of Physiological Chemistry I, Department of Medical Biochemistry and Biophysics, Karolinska Institutet, SE-17 177 Stockholm, Sweden; Department of Neurology, University of Massachusetts, Worcester, MA, USA; Department of Neurology, Beth Israel Deaconess Medical Center, Boston, MA, USA; Department of Chemistry and Chemical Biology, Harvard University, Cambridge, MA, USA; Department of Cell Biology, Harvard Medical School, Boston, MA, USA

**Keywords:** Plasma biomolecules, magnetic levitation, multiple sclerosis, proteomics, lipidomics, metabolomics

## Abstract

We recently discovered that superparamagnetic iron oxide nanoparticles (SPIONs) can levitate plasma biomolecules in the magnetic levitation (MagLev) system and cause formation of ellipsoidal biomolecular bands. To better understand the composition of the levitated biomolecules in various bands, we comprehensively characterized them by multi-omics analyses. To probe whether the biomolecular composition of the levitated ellipsoidal bands correlates with the health of plasma donors, we used plasma from individuals who had various types of multiple sclerosis (MS), as a model disease with significant clinical importance. Our findings reveal that, while the composition of proteins does not show much variability, there are significant differences in the lipidome and metabolome profiles of each magnetically levitated ellipsoidal band. By comparing the lipidome and metabolome compositions of various plasma samples, we found that the levitated biomolecular ellipsoidal bands do contain information on the health status of the plasma donors. More specifically, we demonstrate that there are particular lipids and metabolites in various layers of each specific plasma pattern that significantly contribute to the discrimination of different MS subtypes, i.e., relapsing-remitting MS (RRMS), secondary-progressive MS (SPMS), and primary-progressive MS (PPMS). These findings will pave the way for utilization of MagLev of biomolecules in biomarker discovery and diagnosis of this and other complex disorders.

## 1. Introduction

Magnetic levitation of diamagnetic materials/objects in paramagnetic solutions is a well-documented density-based analysis and separation technique that has been used for a wide range of biological applications from cell sorting and tissue engineering to bone regeneration and self-assembly of living materials.(Abrahamsson et al. 2020; Baday et al. 2019; Deshmukh et al. 2021; Durmus et al. 2015; Gao et al. 2022; Ozefe and Arslan Yildiz 2020; Parfenov et al. 2020a; Parfenov et al. 2020b; Puluca et al. 2020; Sözmen and Arslan-Yıldız 2022; Tasoglu et al. 2013; Tasoglu et al. 2015; Tocchio et al. 2018; Yang et al. 2019) However, this technique cannot levitate nanoscale biomolecules (e.g., plasma proteins), at least in part, due to i) the Brownian motion effect, and ii) the instability of the biomolecules arising from their interactions with high concentrations of conventional paramagnetic salts (e.g., GdCl_3_ and MnCl_2_).(Ashkarran and Mahmoudi 2021),(Ashkarran et al. 2020b)

To overcome these limitations, we recently introduced superparamagnetic iron oxide nanoparticles (SPIONs), constituting a novel paramagnetic liquid that is able to minimize Brownian motion and reliably/reproducibly levitate plasma biomolecules. This is due mainly to SPIONs’ higher magnetic susceptibility based on their superparamagnetic properties as compared to paramagnetic salts, and their interactions with biomolecules in terms of inducing instability to their structures. Therefore, the use of SPIONs allows safer and faster levitation of nanoscale plasma biomolecules, compared to conventional paramagnetic solutions. (Ashkarran and Mahmoudi 2021; Ashkarran et al. 2020b) One of our striking observations from using SPIONs was that upon injection of the plasma into the MagLev system, plasma biomolecules began forming several ellipsoidal layers.(Ashkarran et al. 2020a) Another striking finding was that the evolving magnetically levitated plasma biomolecules provided useful information regarding the health of plasma donors.(Ashkarran et al. 2020a) To uncover the mechanism behind the formation of these unique patterns, we conducted proteomics analysis on the ellipsoidal layers but found no remarkable difference in their proteome profiles.(Ashkarran et al. 2020a)

To mechanistically understand the underlying phenomena behind the formation of ellipsoidal bands in the magnetically levitated plasma biomolecules, here we report multi-level omics analysis on the formed ellipsoidal bands using lipidomics and metabolomics analysis. To investigate whether such differences in the biomolecular composition of various levitated ellipsoidal bands actually reflect health information, we conduct a proof-of-concept study on plasma samples of individuals who suffer from various types of multiple sclerosis (MS); we used MS subtypes as model diseases because discriminating among them is a major clinical challenge in autoimmune medicine.

MS is a common, unpredictable, immune-mediated, complex demyelinating disorder that affects the central nervous system (CNS). (Reich et al. 2018; Vaughn et al. 2019; Wootla et al. 2012) There are four main types of MS: relapsing-remitting MS (RRMS), secondary-progressive MS (SPMS), primary-progressive MS (PPMS), and progressive-relapsing MS (PRMS).(Axtell et al. 2010; DeLuca et al. 2020) Currently, clinicians must rely on combining the results of various clinical examinations including magnetic resonance imaging (MRI) and cerebrospinal fluid (CSF) analysis to determine the disease phenotype.(Absinta et al. 2016; Cheng et al. 2017; Vrenken et al. 2010) Therefore, there is serious need for development of new techniques for early identification of different clinical phenotypes and patients at risk for progressive disease in order to accelerate biomarker detection that can help us understand MS pathogenesis, monitor disease progression, prescribe appropriate therapy early on, and contribute to novel therapeutics.(Knowlton et al. 2015; Knowlton et al. 2017)

## 2. Materials and methods

### 2.1. Materials

SPIONs (30 mg/ml, commercially known as ferumoxytol) was purchased from Feraheme (http://www.feraheme.com) and diluted with phosphate-buffered saline (PBS 1X, HyClone) solution to desired concentration for all experiments. Plasma proteins from patients with three different types of MS were provided from the MS center at the university of Massachusetts Medical Center, Worcester, MA. For calibration of the MagLev system, fluorescent polyethylene microparticles with known densities and standard-density solid-glass particles were obtained from Cospheric (http://www.cospheric.com) and American density materials (http://www.americandensitymaterials.com), respectively.

### 2.2 MagLev platform

The standard MagLev platform used in the present work is depicted in **Figure 1**. The standard MagLev set-up is one of the simplest configurations: have two blocks of N42-grade neodymium (NdFeB) cubic magnets (25.4 mm length, 25.4 mm width, and 50.8 mm height), 2.5 cm distance of separation and (like other MagLev platforms) the N poles face each other. Disposable plastic cuvettes were cut to 25mm and used as levitation containers. All common MagLev devices have two blocks of coaxial identical permanent magnets with face-to-face similar poles and a separation distance of “d”. The details of the relationship between the density of an object and its levitation height within the MagLev system are well understood and reported by our group and others.(Alseed et al. 2021; Ge et al. 2020; Turker and Arslan-Yildiz 2018)

**Figure 1:**
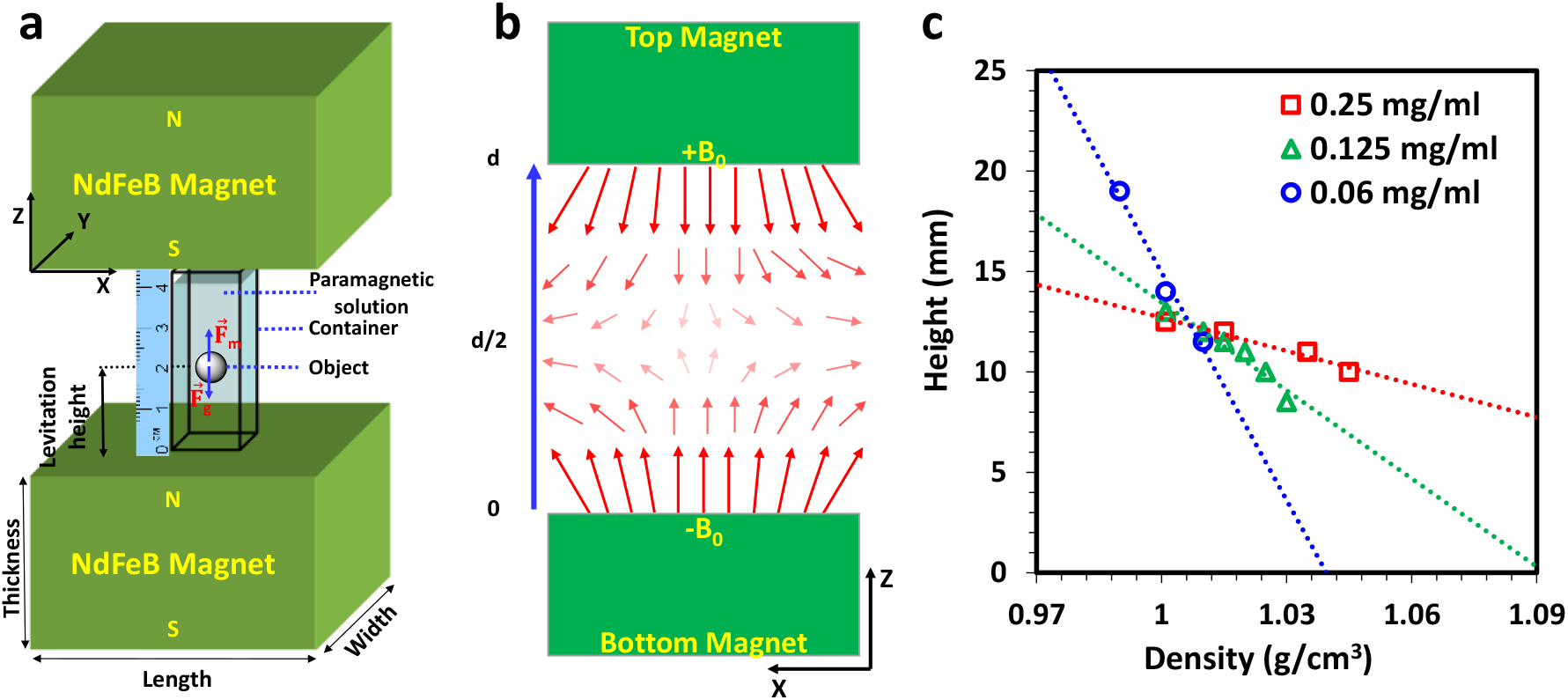
Overall configuration and calibration of the MagLev system. (a) schematic of the standard MagLev configuration used herein, (b) the pattern of the magnetic field between the magnets, and (c) levitation heights of standard density glass beads versus their densities at various concentration of SPIONs showing calibration of the MagLev system.

Briefly, the “density-levitation height” equation can be derived simply by considering the anti-Helmholtz configuration of magnetic fields, which results in a B-field only in the z direction (i.e., B-fields in XY plane cancel out each other):(Ge et al. 2018)

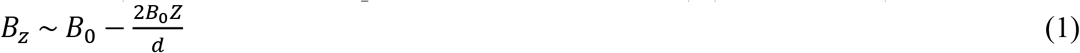

The mechanical equilibrium of the levitating object is achieved due to the balance of gravitational and magnetic forces within the MagLev system, in a 3D Cartesian coordinate system by assuming the z-axis as a symmetry axis of the MagLev:

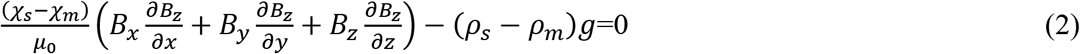

One can simply derive the relation between the density of the object and its levitation height as:

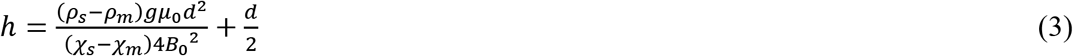

Where, ρ_m_ and ρ_s_ (kg/m^3^) are the density of the paramagnetic medium and sample respectively, g (m/s^2^) is the gravitational acceleration, μ_0_ (T.m.A^−1^) is the permeability of free space, d (m) is the distance between the magnets, B_0_ (tesla) is the magnitude of the magnetic field at the surface of the magnets, χ_m_ and χ_s_ (unitless) are the magnetic susceptibilities of the paramagnetic medium and the sample, respectively.

### 2.3. Characterization

Two blocks of cubic-shaped NdFeB permanent magnets (grade N42, Model # NB044) were obtained from magnet4less. Magnetic field strength (~ 0.5 T on both magnets’ surfaces) between the magnets was measured by a gauss meter (vector/magnitude Gauss meter model VGM, Alphalab). Levitation profiles of the particles were recorded by a Nikon D750 digital camera containing a 105 mm Nikkor Microlens and a millimetre-scale ruler.

### 2.4. Proteomics sample preparation

Sample preparation was performed according to previous reports with slight modifications.(Saei et al. 2018) The protein concentration was measured using Pierce BCA protein assay (Thermo); 10 μg of protein from each sample was transferred to a new tube, and urea was added to a final concentration of 1 M. The extracted proteins were reduced with dithiothreitol (DTT) to a final concentration of 10 mM (37°C for 45 min) and alkylated with 50 mM iodoacetamide (IAA) at room temperature for 45 min in the dark. After precipitation of samples with methanol/chloroform, samples were dissolved in Tris buffer with 8 M urea and diluted to 1 M urea after resuspension. Then, modified sequencing-grade trypsin (Promega) was added at a ratio of 1:50 trypsin to protein followed by overnight incubation. Subsequently, samples were acidified and cleaned using StageTips (Thermo) according to the manufacturer’s protocol. Samples were dried using SpeedVac centrifugal evaporator, and prior to LC-MS/MS analysis, 0.1% formic acid (FA) (+2% acetonitrile (ACN)) solution (Fluka) was added to the samples to achieve a concentration of 0.2 μg/μl.

### 2.5. Proteomics analysis

Protein digests (1 μg) were loaded with buffer A onto a 50-cm column [EASY-Spray, 75 μm internal diameter (ID), PepMap C18, 2 μm beads, 100 Å pore size] connected to a nanoflow Dionex UltiMate 3000 UHPLC system (Thermo) and eluted in an organic solvent gradient increasing from 4% to 26% (B: 98% ACN, 0.1% FA, 2% H_2_O) at a flow rate of 300 nL/min over 80 min.

Mass spectra were acquired with an Orbitrap Fusion mass spectrometer (Thermo Fisher Scientific) in data-dependent mode at MS1 resolution of 120,000 and MS2 resolution of 30,000, in the m/z range from 350 to 1800. Peptide fragmentation was performed via higher-energy collision dissociation (HCD) with energy set at 30 NCE.

### 2.6. Proteomics data processing

The raw data from mass spectrometry were analyzed by MaxQuant, version 1.5.6.5.(Cox and Mann 2008) The Andromeda search engine searched MS/MS data against the International Protein Index database (human version 2014_02, 89,054 entries).(Cox et al. 2011) Mass tolerance for precursor ions was 20 ppm (initial search) and 4.5 ppm (main search), and the MS/MS mass tolerance was set at 20 ppm. Cysteine carbamidomethylation was used as a fixed modification, whereas methionine oxidation was selected as the variable modification. Trypsin/P was selected as the protease and no more than two missed cleavages were allowed. A 1% false discovery rate was used as a filter at both protein and peptide levels. For all other parameters, the default settings were used. Label-free quantification of peptides and proteins was performed. Only proteins with at least two peptides were included in the final data set, and all contaminants were removed.

### 2.7. Lipidomics and metabolomics sample preparation, analysis and data processing Sample extraction

Maglev fractions on ice were spiked with synthetic lipid and polar metabolite internal standards (500 total picomoles) of di-myristoyl phospholipids (PC, PG, PE, PS, PA), 50 picomoles SM (30:1) and 25 picomoles TG (14:1), along with 20 nanograms of D4-glycochenodeoxycholic acid (D4-GCDCA) and 20 nanograms of 1-lyso-myristoyl PC. To each sample, 300 microliters of −20°C chilled methanol containing 1 mM BHT (an antioxidant) was added, and samples were vortexed vigorously for 5 min. One mL of tert-methyl-butyl ether (MTBE) was added to each sample, and samples were vortexed for 60 min at room temperature. 150 μl of water were added, and the samples were vortexed for an additional 15 min and then centrifuged for 15 min. The lipid-containing organic supernatants were collected to new test tubes and precipitated proteins were re-extracted as above. Pooled organic extracts were dried overnight in a speedvac, and resuspended in 100 μl of isopropanol. The lower aqueous phase containing polar metabolites was collected to a new tube, dried under vacuum, and resuspended in 100 μl of 95% acetonitrile.

### 2.8. Instrument parameters for global lipidomics analysis

Immediately prior to analysis, aliquots of each lipid extract were diluted in isopropanol:methanol (2:1, v:v) containing 20 mM ammonium formate. Full scan MS spectra at 100,000 resolution (defined at m/z 400) were collected on a Thermo Scientific LTQ-Orbitrap Velos mass spectrometer in both positive and negative ionization modes. Scans were collected from m/z 200 to m/z 1200. For each analysis, 10 μl of sample was directly introduced by flow injection (no LC column) at 10 μl/min using a heated electrospray ionization (HESI) source. A Shimadzu Prominance HPLC served as the sample delivery unit. The sample and injection solvent were 2:1 (v: v) isopropanol: methanol containing 20 mM ammonium formate. The spray voltage was 4.5 kV, ion transfer tube temperature was 275 °C, the S-lens value was 50 percent, and the ion trap fill time was 100 ms. The autosampler was set to 15 °C. After two min of MS signal averaging, the LC tubing, autosampler, and ESI source were flushed with 1 mL of isopropanol, prior to injection of the next sample. Samples were analyzed in random order, interspersed by solvent blank injections, extraction blank injections, and pooled QC samples derived from all study samples. Following MS data acquisition, offline mass recalibration was performed with the “Recalibrate Offline” tool in Thermo Xcalibur software according to the vendor’s instructions, using the theoretical computed masses for the internal calibration standards and several common endogenous mammalian lipid species. MS/MS confirmation and structural analysis of lipid species identified by database searching were performed using higher-energy collisional dissociation (HCD) MS/MS at 60,000 resolution and a normalized collision energy of 25 for positive ion mode, and 60 for negative ion mode. MS/MS scans were triggered by inclusion lists generated separately for positive and negative ionization modes.

### 2.9. Lipid peak finding, identification, and quantification

Lipids were identified using the Lipid Mass Spectrum Analysis (LIMSA) v.1.0 software linear fit algorithm, in conjunction with an in-house database of hypothetical lipid compounds, for automated peak finding and correction of ^13^C isotope effects. Peak areas of found peaks were quantified by normalization against an internal standard of a similar lipid class. The top ~300 most abundant peaks in both positive and negative ionization mode were then selected for MS/MS inclusion lists and imported into Xcalibur software for structural analysis as described above. For this untargeted analysis, no attempt was made to correct for differences in lipid species ionization due to the length or degree of unsaturation of the esterified fatty acids. Therefore, lipid abundance values are inherently estimates rather than true ‘absolute’ values.

### 2.10. QA/QC measures

Prior to the analysis, the mass spectrometer inlet capillary was removed and cleaned by sonication, and the ESI tubing and spray needle were cleaned by extensive flushing with isopropanol. During the analysis, several injection solvent blanks and extraction blanks were interspersed between the randomized study samples to monitor for background ions and any potential sample carryover throughout the run. A pooled quality control sample was created using an aliquot of each study sample, and the pooled QC was injected at the beginning, in the middle, and at the end of the instrument run to monitor for potential drift in instrument response and ensure in-run reproducibility of measurements. The offline external mass recalibration routine described above was used to eliminate in-run drift in mass calibration and improve the consistency of mass accuracy over the measured m/z range.

Each lipid species is identified at minimum as a sum composition of lipid headgroup and fatty acyl total carbons: total double bonds, based on accurate mass measurement and isotopic distribution.

Under the extraction conditions used, some lipid species are partially or completely lost during phase separation and therefore are often not reliably quantified with this method. These include free sphingosine and its analogues and derivatives, as well as gangliosides and other highly polar lipid species.

### 2.11. LC gradient for global metabolomics

The LC-MS platform consisted of a Shimadzu Prominence HPLC coupled to a Thermo LTQ-Orbitrap Velos mass spectrometer. The LC system included two LC20AD pumps, a vacuum degassing system, autosampler, and column oven. The HPLC column was a Phenomenex 2.0 mmx100 mm HILIC (3 micron, 100 Angstrom pore size) column equipped with a guard cartridge of the same column chemistry. Solvent A was water containing 50 mM ammonium formate. Solvent B was acetonitrile. The flow rate was 200 μl per min and the column oven was held at 25 °C. The autosampler was held at 4 degrees C. 10 μl of each sample was injected per analysis. The gradient conditions used were: Time 0-2 minutes, 95% solvent B. Column eluant was diverted to waste using a 2-position 6 port valve for the first 1.5 min. From time =2.0 min to 14.0 min, Solvent B was decreased linearly to 50%, then held at 50% for 5 min. The column was then requilibrated at 95% B for 10 min.

### 2.12. Mass spectrometry for global metabolomics

Column eluent was introduced to a Thermo LTQ-Orbitrap Velos mass spectrometer via a heated electrospray ionization source. The mass spectrometer was operated in negative ion mode and positive ion modes separately for separate analytical runs of each sample. Instrument resolution was set to 60,000 using the FT analyzer for full scan MS data, and data-dependant product ion spectra were collected on the 4 most abundant ions at 7,500 resolution using the FT analyzer. The electrospray ionization source was maintained at a spray voltage of 4.5kV with sheath gas at 30 (arbitrary units) and auxiliary gas at 10 (arbitrary units). The inlet of the mass spectrometer was held at 350 °C, and the S-lens was set to 35%. The heated ESI source was maintained at 350 °C.

### 2.13. Data Analysis for global metabolomics

Chromatographic alignment, isotope correction, peak identification and peak area calculations were performed using MAVEN and XCMS software. Peak areas were normalized against the D4-GCDCA and LysoPC(14:0) internal standard. Analyte m/z and RT were compared against reference standards using MAVEN software, and unknown LC-MS peaks generated by XCMS software were searched for potential matches against the KEGG online database.

Lipid class abbreviations used:

PA=phosphatidic acid
PC=phosphatidyl choline
PE=phosphatidyl ethanolamine
PG= phosphatidyl glycerol
PI= phosphatidyl inositol
PS= phosphatidyl serine
CL= cardiolipin
AC=acylcarnitine
hAC=hydroxy acylcarnitine
Cer= ceramide
HexCer= hexosylceramide
LacCer= Lactosylceramide
CerPO4= ceramide-1-phosphate
ST= sulfatide
SM= sphingomyelin
Sph= sphingosine or sphingosine analogue
Chol=cholesterol, or chol(FA)= cholesteryl ester
MG= monoacylglyceride
DG= diacylglyceride
TG= triacylglyceride
FA= non-esterified fatty acid
e = ether-linked fatty acid (plasmalogen or O-alkyl ether)
O= O-alkyl-ether linked fatty acid
P= Plasmalogen (vinyl ether)-linked fatty acid

## 3. Results

We have recently shown that complex protein environments (i.e., human plasma proteins) form a unique ellipsoidal pattern in the presence of SPIONs when levitated in the MagLev system. Formation of the plasma pattern starts a few minutes after injection of the proteins into the MagLev and evolves over three hours.(Ashkarran et al. 2020b) Such specific and highly reproducible plasma patterns form for plasma proteins (not single proteins or standard density microspheres; see **Supplementary Figure 1** and **Supplementary Figure 2** for levitation profiles of single proteins and fluorescent polyethylene microspheres) regardless of the type of the human plasma (e.g., healthy or disease related). The multiple ellipsoidal bands and corresponding particular plasma patterns that appear may be related to i) configurational or structural variation of individual proteins, ii) protein-protein interactions that generate lesser or greater density solution constructs, and/or iii) interaction of proteins and other biomolecules (i.e., lipids and metabolites) with the magnetic field of the MagLev system. Due to the different physicochemical properties of human plasma biomolecules (e.g., difference in density, charge, molecular weight, viscosity, hydrophilic/hydrophobic ratio, mechanical properties, and protein-protein interaction), they respond differently to an external magnetic field, creating a ‘‘fingerprint’’ for each individual human plasma.

To understand the mechanism behind the formation of plasma patterns (see **Figure 2** for overall flow of the study), plasma biomolecules of various healthy individuals (n=3) and MS patients with three different types of MS [(i.e., RRMS (n=14), SPMS (n=5), and PPMS (n=3)] were magnetically levitated (**Figure 2**; see **Supplementary Figures 3** to **5** and **Supplementary movies 1** to **3** for details on levitation progress and patterns of various types of MS). The plasma samples (n=25) were first levitated in the MagLev system, where they created *six* ellipsoidal bands of plasma biomolecules, validating the high reproducibility of the patterns of levitated plasma proteins regardless of the source of plasma.

**Figure 2:**
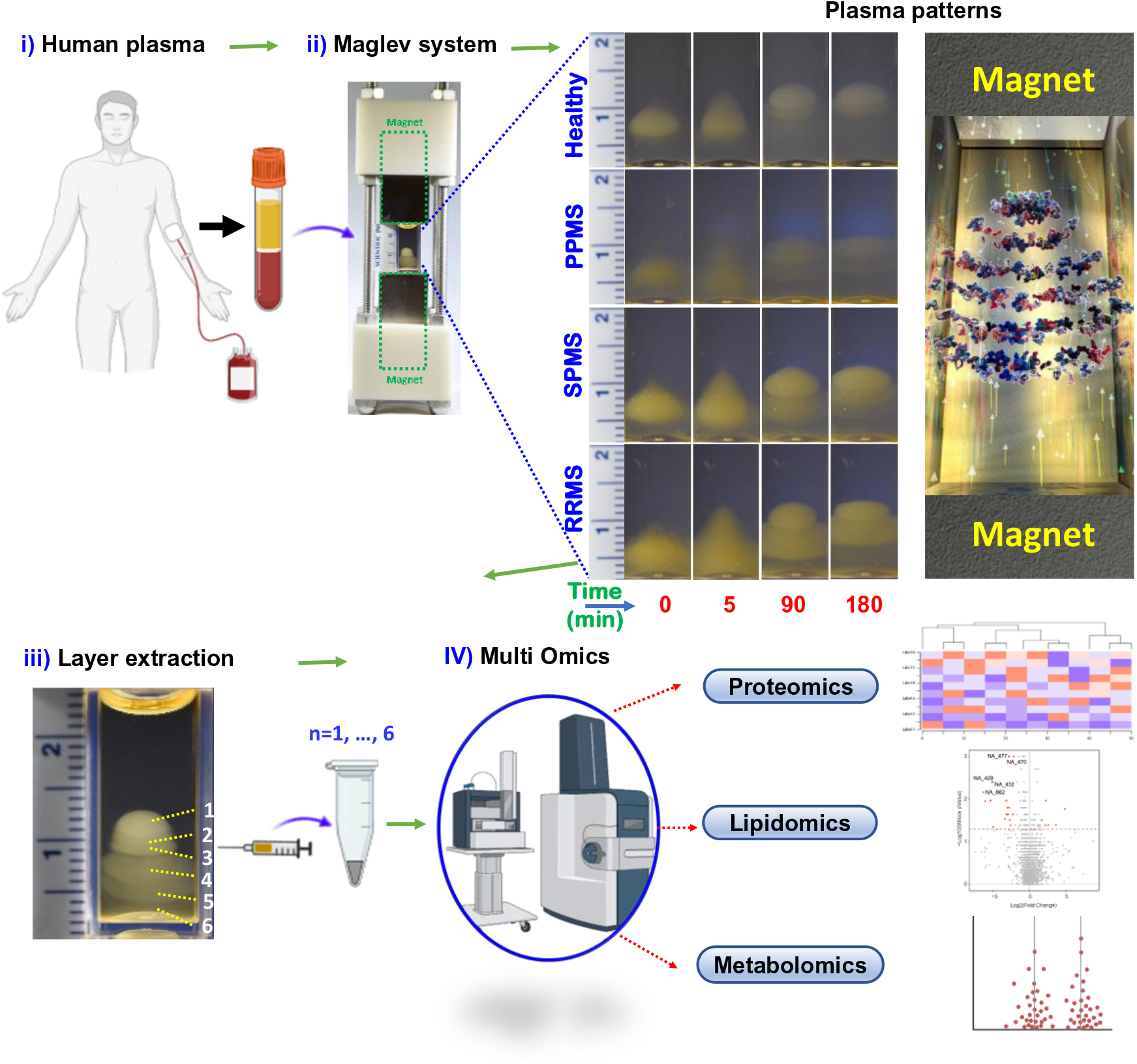
Schematic representation of the study and plasma protein patterns within the MagLev system. Formation of ellipsoidal patterns over time from levitated plasma biomolecules (n=25).

**Figure 2:**
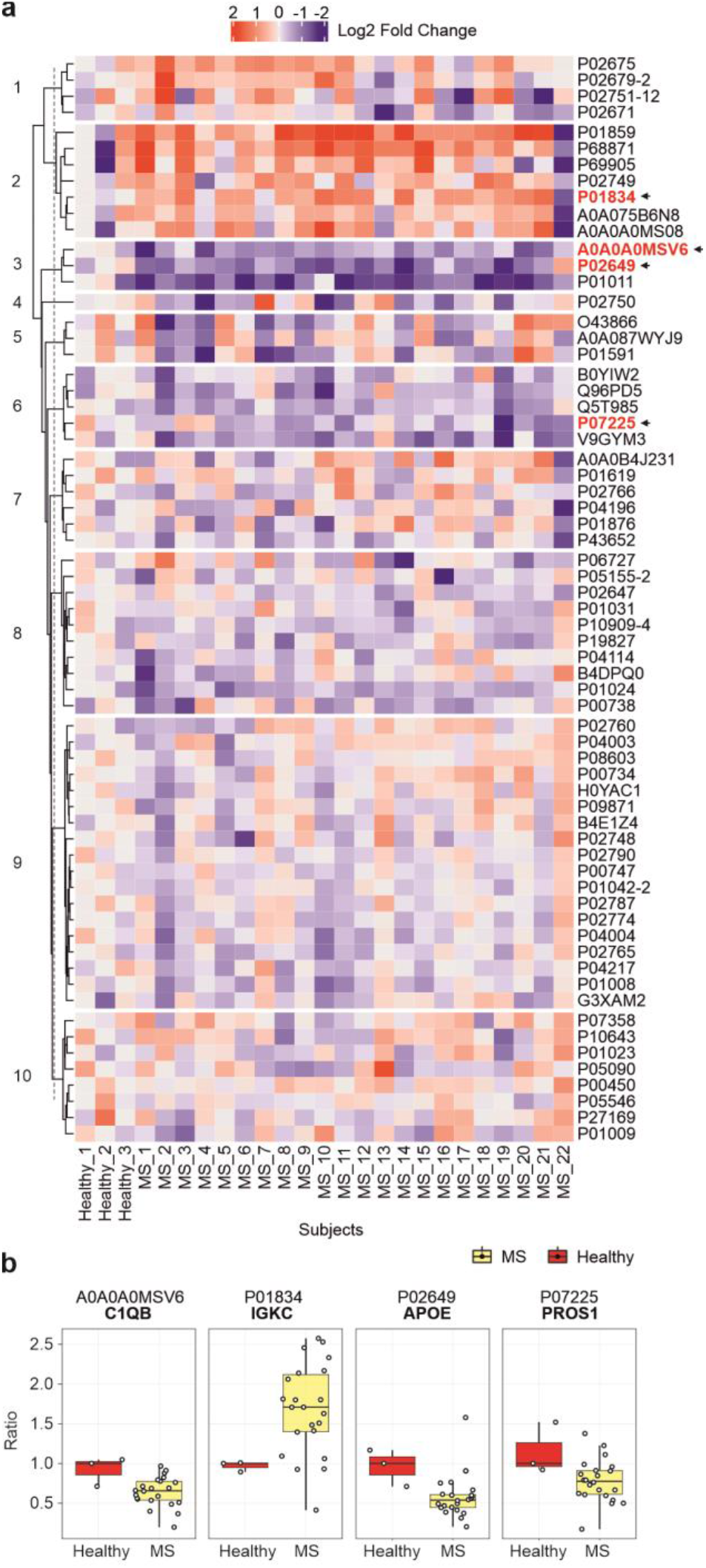
Protein heatmap of MS patients vs. healthy individuals. **(a)** The relative abundance of plasma proteins in healthy subjects vs. patients with different types of MS shows considerably individual variability. **(b)** Four proteins that are significantly different (p < 0.05) in MS subjects vs. healthy (Wilcoxon non-parametric sum test). UniProt names are shown above, and gene names are shown in bold above each boxplot (Center line, median; box limits contain 50%; upper and lower quartiles, 75 and 25%; maximum, greatest value excluding outliers; minimum, least value excluding outliers; outliers, more than 1.5 times of upper and lower quartiles).

To mechanistically explore the driving force that dictates formation of ellipsoidal bands, we extracted the bands layer by layer from top to bottom of the MagLev column and carried out multi-omics analysis of their biomolecular composition to identify proteomic, lipidomic, and metabolomic profiles in all 25 plasma samples.

### 3.1. Proteomics

We analyzed the bulk plasma proteomes of the 25 plasma samples (3 healthy and 22 MS patients). Quantifying 210 proteins with at least two peptides for each protein, the data were normalized by the sum of all the protein intensities in each sample (**Supplementary Data 1**). For analysis of various layers in each individual MS sample, 3 healthy samples were used as control and compared with 22 MS patients. For each protein, the abundance was divided by the median intensity of the 3 healthy samples, and the determined value (fold change) was log2 transformed. The corresponding heatmap, presented in **Figure 2a**, represents the 65 proteins quantified across all 25 plasma samples (see **Supplementary Data 1** for details of all detected proteins).

Wilcoxon non-parametric sum test was used for testing significance, comparing fold change in 3 healthy samples *vs*. 22 MS patient samples. Four proteins passed the 0.05 threshold for *p* value, and the relative abundance of these proteins in all MS *vs*. healthy samples is shown in boxplots in **Figure 2b**. Volcano plots for each MS patient *vs*. control (highlighting aforementioned four proteins) are shown in **Supplementary Figure 6**. Three of these proteins are C1QB (Grewal et al. 1999), IGKC (Torkildsen et al. 2010), and APOE,(Burwick et al. 2006) which have previously been associated with MS pathophysiology, suggesting the validity of our outcomes. However, none of these proteins has previously been identified as a MS biomarker. The last protein was PROS1, and its connection to MS has not been shown before. It is noteworthy that we have previously observed there is no significance difference among proteomics profile of various layers of the produced plasma patterns in the MagLev system (Ashkarran et al. 2020a).

### 3.2. Lipidomics

To further investigate the effect of the MagLev system on other biomolecules, we conducted lipidomic analysis. Twenty-four plasma samples (3 healthy and 21 MS individuals) were subjected to MagLev, and 6 different layers of the emerging plasma patterns of each sample (144 layer in total) were analyzed (one of the samples was removed due to technical issues), and 312 lipid species were reliably identified and quantified across the samples. The raw intensities were normalized by the sum of all lipid intensities in each sample (**Supplementary Data 2)**. Further normalization was performed to adjust the median intensity of lipids to 1 (log2 of 0). For each lipid, abundance was divided by the median intensity of the corresponding layers in 3 healthy samples, and the determined value (fold change) was log2 transformed. The corresponding heatmap depicted in **Figure 3** shows the overall lipid intensities across all 144 samples (protein cluster assignment information in **Supplementary Data 3**). Generally, the lipid profiles show greater variability among individuals compared to the proteomics profiles. While many lipid species are presented uniformly across all samples, lipids in clusters 4, 9, and 10 show great variability between individuals. Therefore, we can deduce that lipids are partially responsible for the differential MagLev patterns of different MS types vs. healthy individuals.

**Figure 3.**
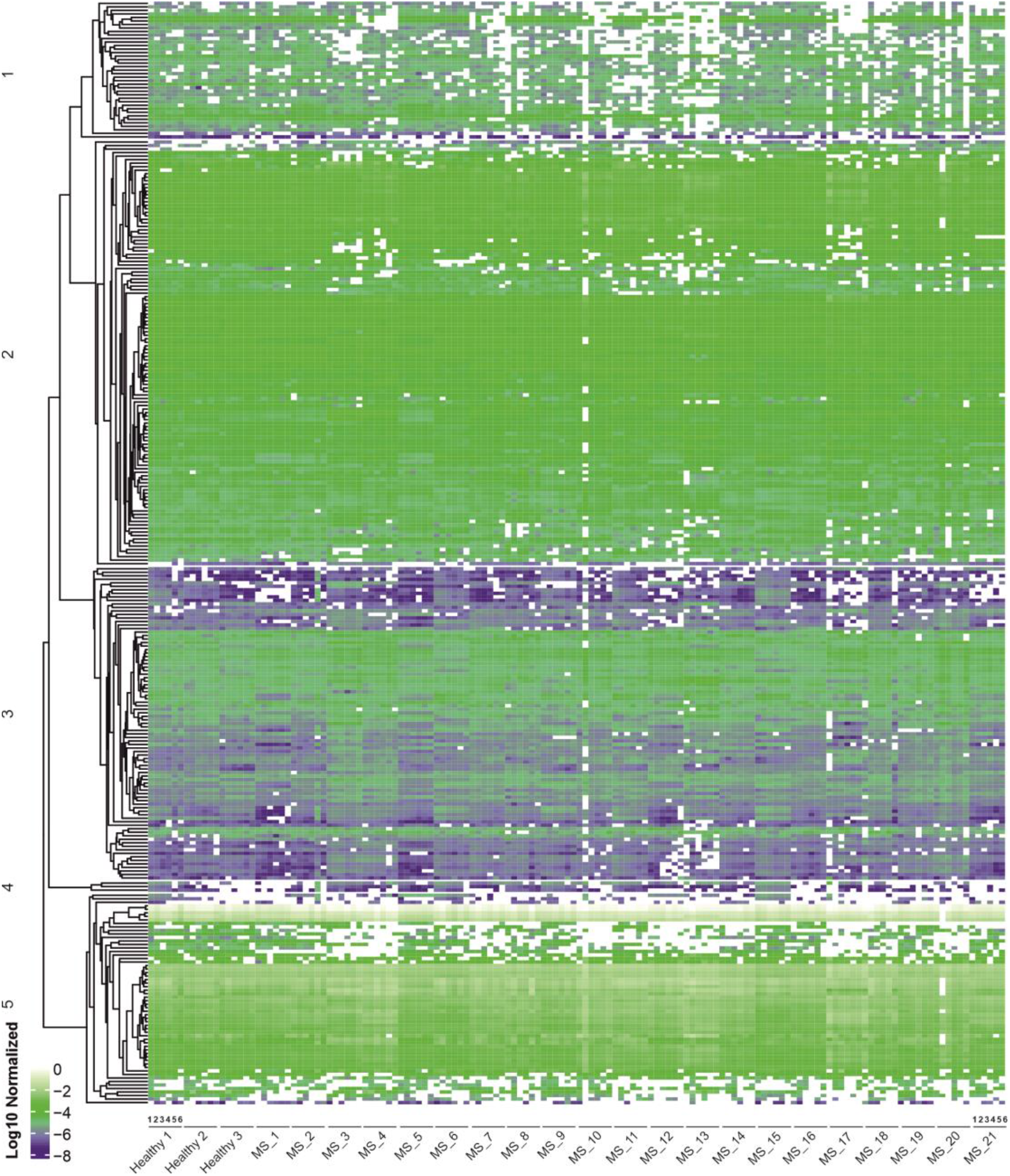
Heatmap of plasma lipidomics profiles for MS patients vs. healthy subjects. The relative abundance of plasma lipids in healthy subjects vs. patients with different types of MS shows both the consistency of some lipid features but also great individual variability among the subjects (layers 1 to 6 are shown from left to right for each subject).

We performed a principal component analysis (PCA) of normalized lipid intensities across all the samples and the results are shown in **Figure 4a**. The first PCA component could separate the RRMS samples from others to some extent. The lipids that made the greatest contribution to the separation of samples along the first and second PCA components are highlighted in the PCA loading scatter plot (**Figure 4b**).

**Figure 4.**
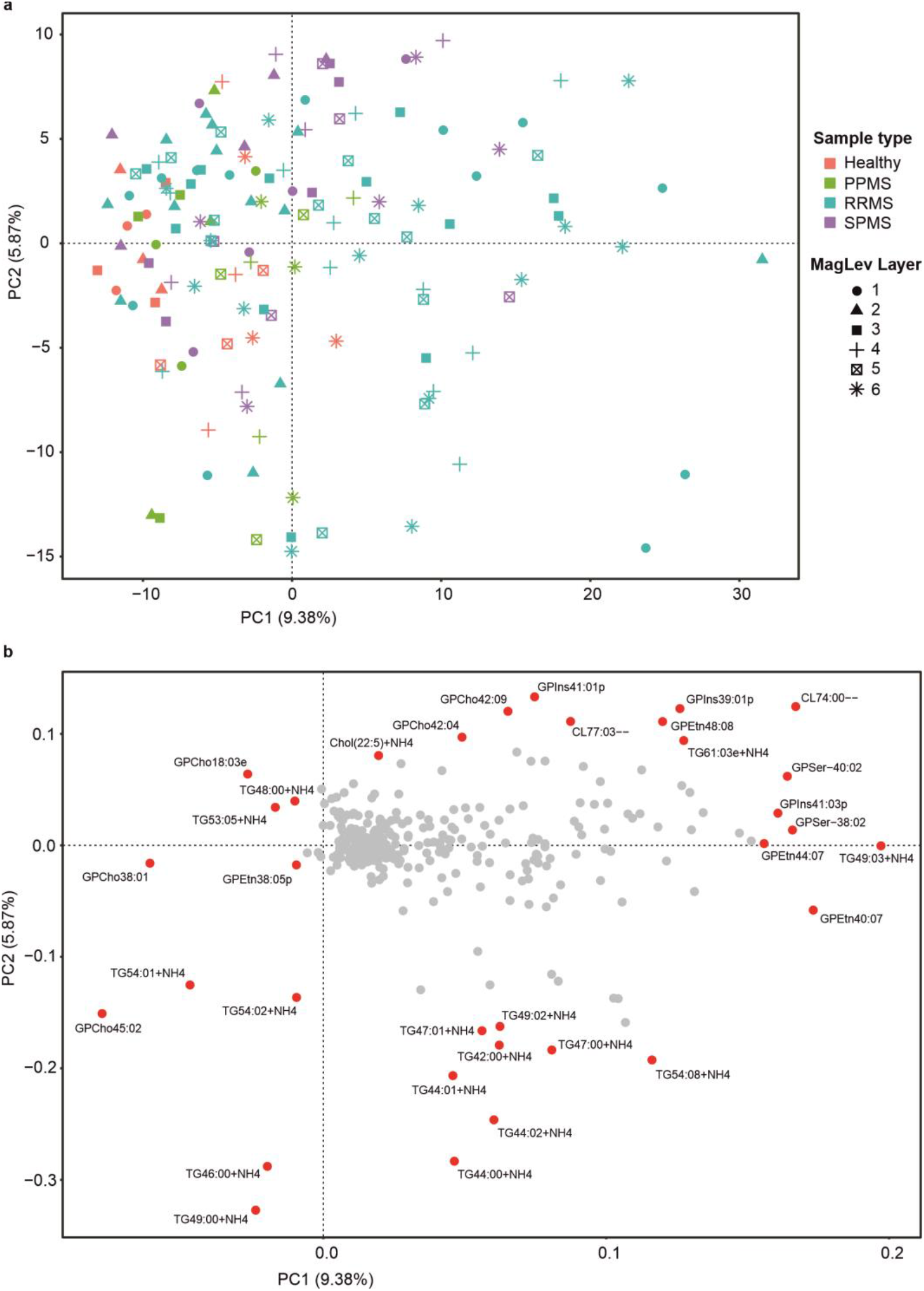
Exploration of the lipidomics data of MagLev layers for MS vs. healthy samples. **(a)** PCA analysis of lipid profiles separates the RRMS samples from other MS types and healthy controls to some extent. But the MS types or different layers are not separated, indicating great variability among subjects. Different colors represent different MS types, and the shapes represent various layers. **(b)** The lipid species that contribute most to the separation of samples along the first and second principal components are highlighted in the PCA loading scatter plot.

Wilcoxon non-parametric sum test was used to test significance, comparing the fold change of each lipid species in 3 healthy samples vs. 21 MS patient samples in each individual layer; results are shown for individual layers in **Figure 5**, highlighting the significant outliers. Significant outliers among lipid species can be noted in most layers, especially in layers 1 and 3-6. Most of the outliers are unique to a specific layer.

**Figure 5.**
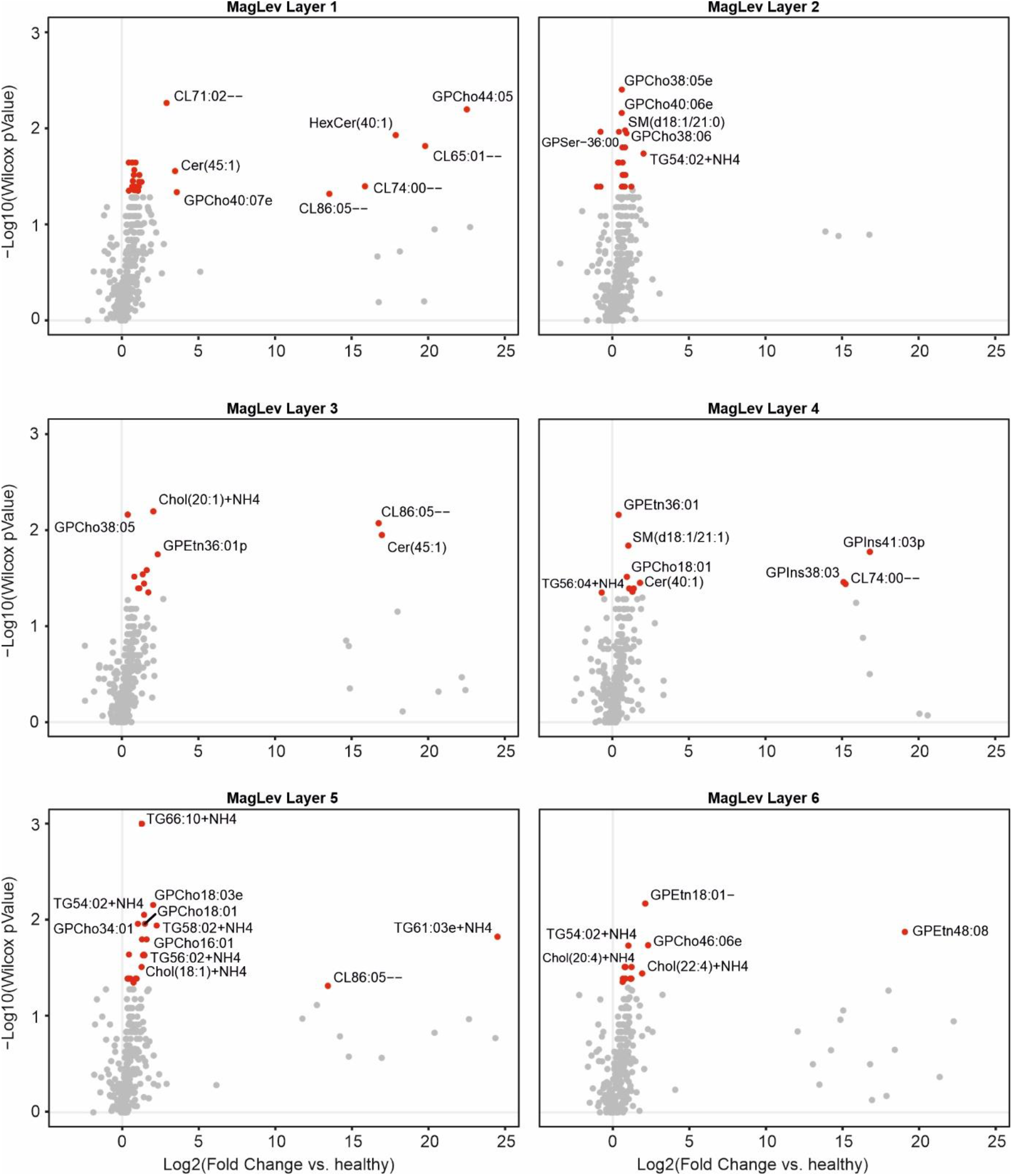
The relative abundance of lipids in MagLev layers of MS patients vs. healthy subjects. The lipid species that differ significantly between MS patients and healthy individuals in each MagLev layer (Wilcoxon *p* value < 0.05).

A number of the identified lipids including glycerophosphochcoline (GPCho) and LysoPC (Boulanger et al. 2000; Del Boccio et al. 2011) have been previously associated with MS pathophysiology, indicating the validity of our analyses. However, none of these lipids have previously been established as MS biomarkers.

Furthermore, we analyzed the data for each layer separately for each MS type and presented the results in volcano plots (**Figure Supplementary Figure 7** and **Supplementary Data 4**). While several lipid species differed significantly in different MagLev layers for SPMS and RRMS, there were no significant outliers in PPMS plots (therefore not shown).

In addition to PCA, we performed PLS analysis (models were built for each layer separately) to investigate whether the model can distinguish healthy individuals from MS patients based on the lipidomics data in each layer. We found that PLS models of all layers could separate healthy from MS samples (see the details of the obtained results for each individual layer from layer 1 to layer 6 in **Supplementary Figures 8** and **9**). Panel “**a**” of the figures shows the PLS model, while panel “**b**” shows the contribution of specific lipids separating the samples. The lipids on the x extremities of panel B are those contributing most to the separation between healthy and MS samples.

Moreover, we investigated whether the PLS models can also discriminate various types of MS based on the lipidomics data from each individual layer separately. Details of the results from PLS models and their loadings are depicted in **Supplementary Figures 10** and **11**). Among the different MS types, RRMS is better separated from the other MS types, and only layer 5 data clearly separated all the MS types. Therefore, with regards to lipidomics analysis, MagLev layer 5 might be of higher diagnostic value in MS biomarker discovery, which can be potentially further improved by a larger cohort size. In the PLS model for layer 1, SPMS and PPMS are cleanly separated from RRMS in the first component, and then PPMS is separated from the other MS types along the second component (**Supplementary Figure 10a**).

### 3.3. Metabolomics

We next performed metabolomics analysis on various layers of the plasma patterns of 24 subjects (3 healthy and 21 MS individuals) in the MagLev system. Similarly, 6 different layers of the plasma patterns for each sample (144 layers in total) were separated and analyzed (similar to the lipidomics analysis, one of the MS samples was removed due to technical issues). 4,343 metabolite species were quantified across the samples, with 382 metabolites identified. Raw intensities were normalized by the sum of all metabolite intensities in each sample (**Supplementary Data 5**). Further normalization was performed to adjust the median intensity of metabolites to 1 (log2 of 0). For each metabolite, abundance was divided by the median intensity of the corresponding layers in 3 healthy samples, the determined value (fold change) was log2 transformed, and the corresponding heatmap (**Figure 6)** shows the overall metabolite intensities across all 144 samples (cluster assignment information presented in **Supplementary Data 6**). While many metabolites are presented uniformly across samples, metabolites in clusters 4, 9, and 10 show great variability between individuals. Therefore, similar to lipids, metabolites are also partially responsible for the differential MagLev patterns of different MS types vs. healthy individuals.

**Figure 6.**
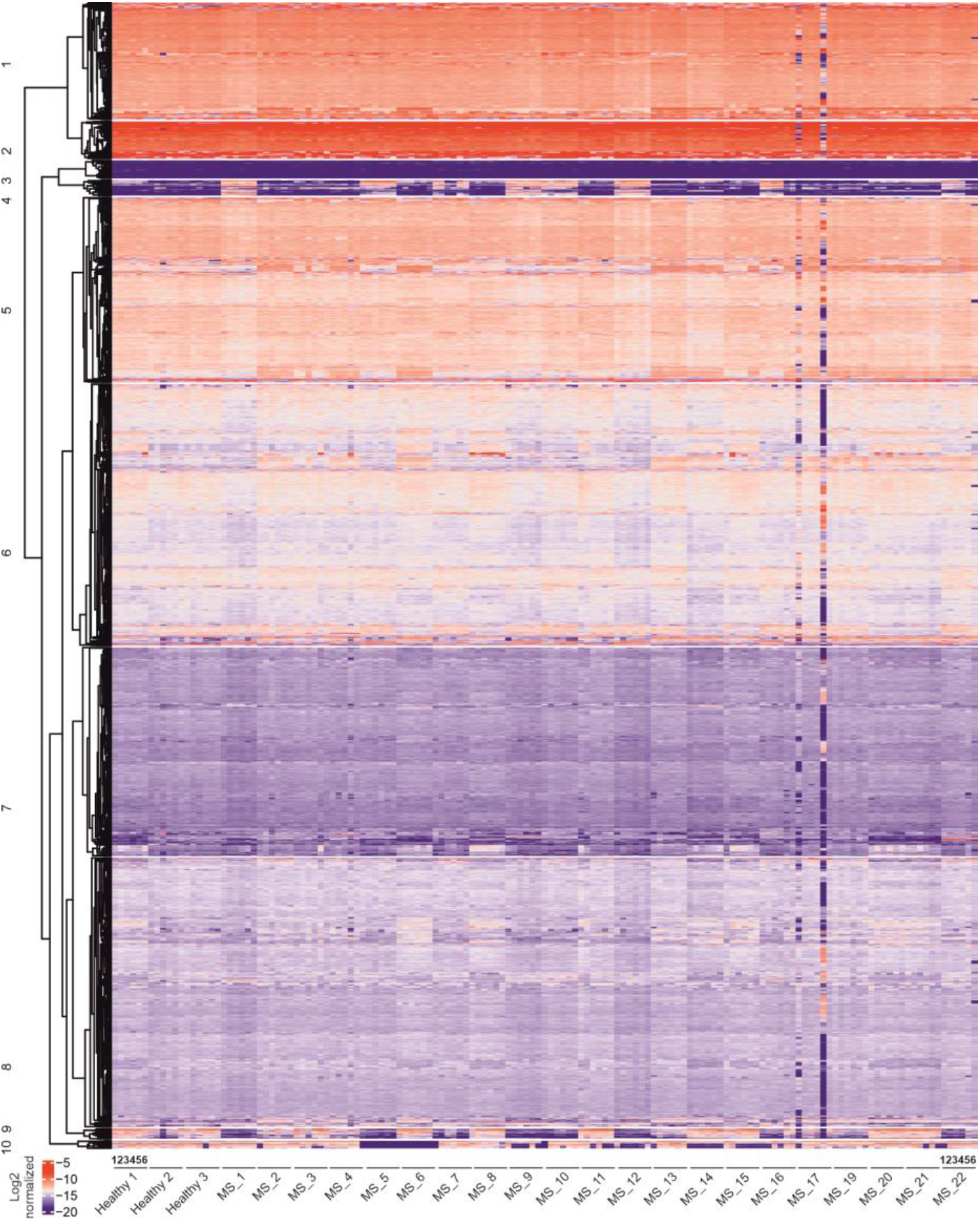
Heatmaps of plasma metabolome profiles for MS patients and healthy subjects. The relative abundance of plasma metabolites in MagLev layers of plasms from healthy subjects vs. patients with different types of MS shows the consistency of some metabolite features but also individual variability among subjects (layers 1 to 6 are shown from left to right for each subject).

A PCA was performed on the normalized metabolite intensities across all the samples, and the results are shown in **Figure 7a**. The first PCA component distinguished healthy samples from PPMS in the majority of cases. Furthermore, the healthy samples could also be differentiated from RRMS to some extent. The metabolomics profiles of the healthy and SPMS, however, were quite similar. The metabolites that contribute most to the separation of samples along the first and second PCs are highlighted in the PCA loading scatter plot (**Figure 7b**).

**Figure 7.**
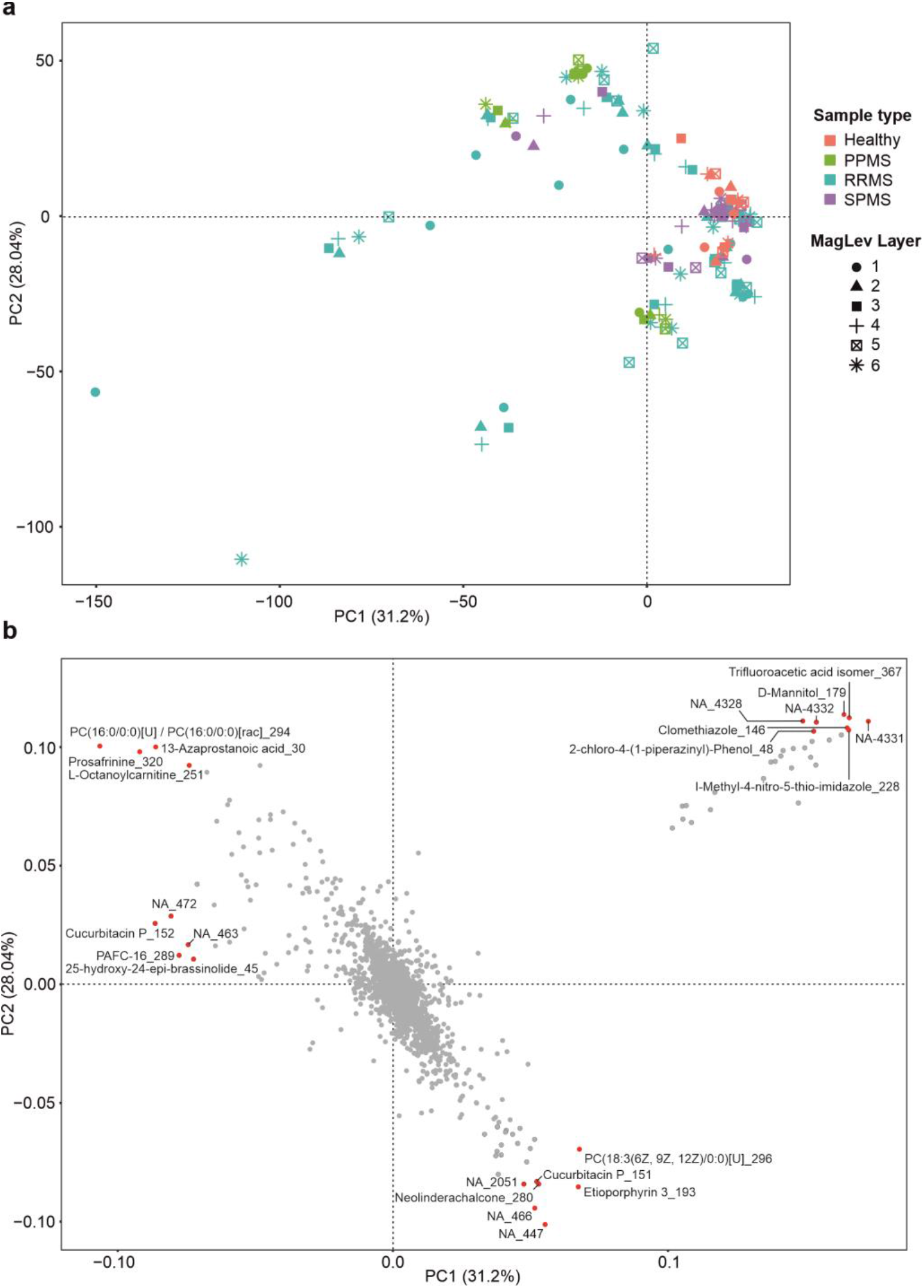
PCA analysis of plasma metabolite profiles show discrimination between samples of healthy origin vs. certain MS types. **(a)** PCA analysis of the samples shows that the plasma metabolite profile can distinguish healthy individuals from MS patients (especially PPMS) to some extent but the MS types or layers are not separated, indicating great variability among subjects, and **(b)** The metabolomes that contribute most to the separation of samples are highlighted in the PCA loading scatter plot. Unidentified metabolites are denoted with NA. The corresponding metabolite IDs are shown after an underscore “_” and can be found in corresponding Supplementary Data files.

Wilcoxon non-parametric sum test was used for testing significance, comparing the fold change of each metabolite in 3 healthy samples vs. 21 MS patient samples in each individual layer, and the results are shown for individual layers in **Figure 8**, highlighting the significant outliers.

**Figure 8.**
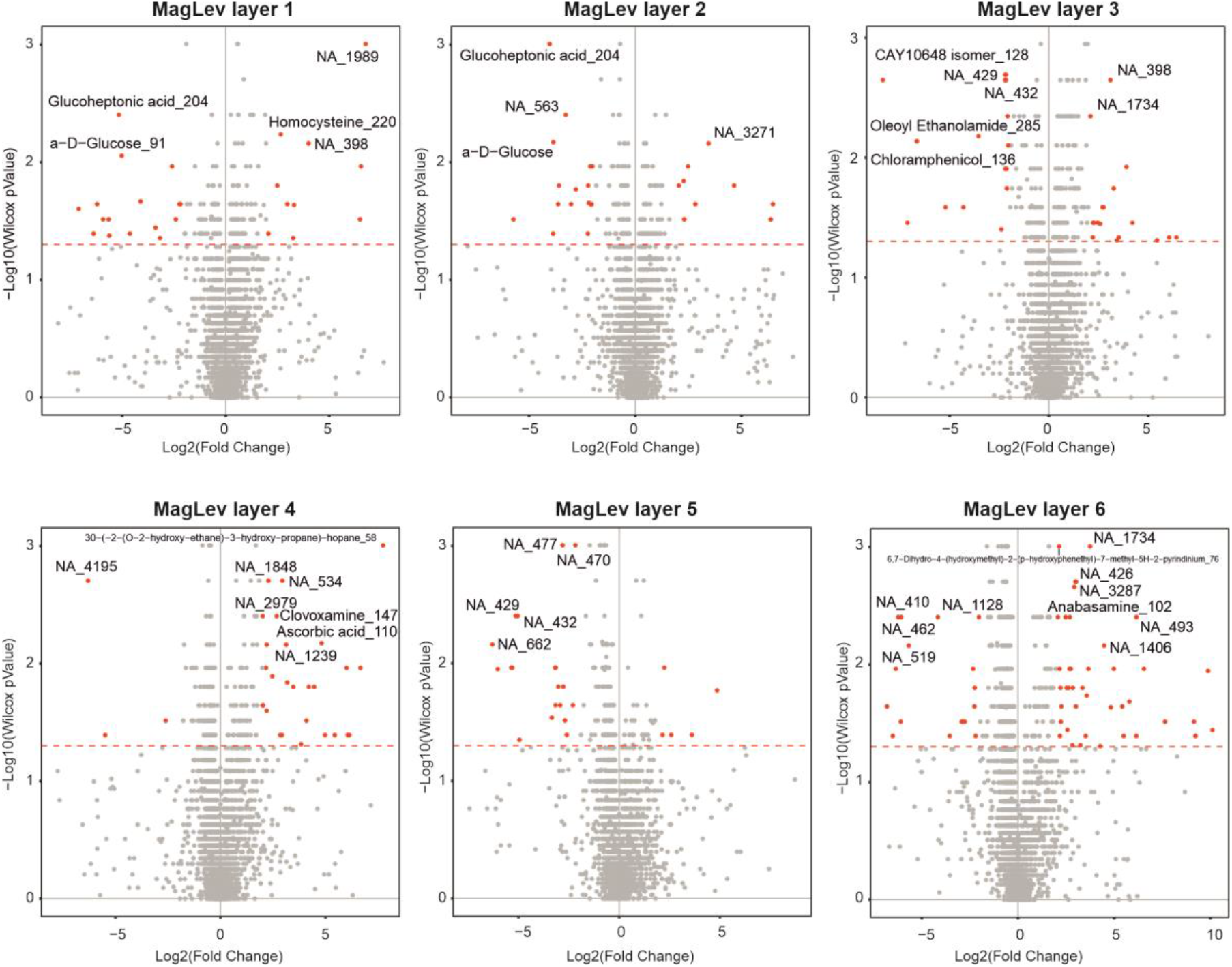
The relative abundance of metabolites in MagLev layers of MS patients vs. healthy individuals. The top metabolites that are significantly different between MS patients and healthy individuals are highlighted in each layer (*p* value < 0.05). The majority of significant outliers are unique to individual layers. The metabolites not identified are denoted with NA. The corresponding metabolite IDs are shown after an underscore “_” and can be found in corresponding Supplementary Data files.

A number of these metabolites including homocysteine and glucoheptonic acid have been previously associated with MS pathophysiology, indicating the validity of our analyses.(Mititelu et al. 2006) However, none of these metabolites has previously been established as an MS biomarker.

Furthermore, we analyzed the data for each layer separately for each MS type, and the results are presented in volcano plots (**Supplementary Figure 12** and **Supplementary Data 7**). Similar to the lipidomics analysis, while several lipid species were significantly altered in different MagLev layers for SPMS and RRMS, there were no significant outliers in PPMS plots (not shown). This indicates that the lipid and metabolite profiles of SPMS and RRMS are very distinct from those of normal patients.

In addition to PCA, we performed PLS analysis (models were built for each layer separately) to investigate whether the model can separate healthy individuals from MS patients based on metabolite profiles. PLS models of all layers were able to distinguish healthy from MS samples (see details for each individual layer from layer 1 to layer 6 in **Supplementary Figures 13** and **14**). Panel “**a**” of the figures shows the PLS model, while panel “**b**” shows the contribution of specific metabolites separating the samples. The metabolites on the x extremities of the panels “**b**” are those contributing most to the separation. Moreover, we showed that the PLS models can also discriminate various types of MS based on the metabolomics data from each individual layer separately (**Supplementary Figures 15** and **16**). Based on component 1 and 2 contributions, layers 1 and 2 could best discriminate the MS types compared to data in the other layers. Therefore, these MagLev layers might have a higher diagnostic value in biomarker discovery for MS in larger cohorts.

## 4. Discussion

We applied multi-level omics analyses to mechanistically understand the reason for formation of various ellipsoidal bands when levitating plasma biomolecules in the MagLev system. Our findings suggest that the significant variations in composition of lipids and metabolomes are the main driving force for formation of various ellipsoidal bands. In addition, we probed multi-omics information across various plasmas to determine whether differences in ellipsoidal patterns hold potential for disease diagnosis and biomarker discovery.

Applying proteomics, lipidomics, and metabolomics to each individual layer of the plasma patterns from healthy subjects and MS patients, we show that lipidome and metabolome profiles (but not proteome profiles) of human plasma are the major determinants of the produced patterns and have significant diagnostic potential.(Del Boccio et al. 2016) While the majority of lipidome and metabolome profiles present in different layers have discriminatory power in biomarker discovery, we demonstrate in this proof-of-concept study that there are particular biomolecules in certain layers of the plasma patterns that have higher diagnostic potential. Although biomarker discovery was not the main focus of the current study, our findings suggest several biomolecules in the plasma patterns that merit further study for discovering novel biomarkers for MS.

Proteomics analysis of various layers of the plasma patterns originating from the MagLev system revealed no significant differences between healthy subjects and MS patients, nor did it fully distinguish different MS subtypes. However, bulk analysis of the plasma samples identified a number of proteins that were significantly different between healthy controls and MS patients. There is cumulative evidence of the involvement of these interesting proteins in neurodegeneration and neuroinflammation.(Han et al. 2008) For instance, C1qb plays diverse neuroprotective roles against pathogens and inflammations in the CNS, and levels of C1qb may be disease specific.(Cho 2019) IGKC and APOE are other critical proteins that are significantly altered (independent of age and sex) when compared with controls in amyotrophic lateral sclerosis (ALS) disease. (Katzeff et al. 2020)

There is growing evidence that changes in the levels of plasma lipids/metabolites may result in neurodegenerative diseases.(Di Paolo and Kim 2011) Lipids and their metabolic products play a critical role in the brain, and aberrations in lipid profiles increases the risk of many types of neurodegenerative disorders, including MS. However, to gain a better understanding of the possible role of various biomolecules in the discrimination of MS subtypes, we need to study larger cohorts, due to the large variability among patients with regards to age, sex, disease type, drugs, weight, BMI, etc. (see demographics information in the SI). Nevertheless, it is well documented that many neurodegenerative diseases are related to pathology and accumulation of disordered lipids and metabolites, which causes toxicity to nerve cells and neurodegeneration.(Di Paolo and Kim 2011; Ferreira et al. 2020) The lipidomics profile of various layers of MagLev patterns reveals the critical role of particular categories of lipids in various layers of the produced patterns, including glyceroPhosphoCholine (GPC), posphatidylcholines (PC), glycerophosphoethanolamine (GPEtn), sphingomyelins (SM), and triacylglycerols (TG). Our results suggest that there is a specific combination of different lipid categories in each individual layer of the produced plasma patterns that may serve as a “fingerprint” for detection/discrimination of various diseases.(Higgs 2010)

Metabolomics is an emerging approach in life science and precision medicine for biomarker discovery in various neurodegenerative diseases such as MS.(Bhargava and Calabresi 2016; Noga et al. 2012) Previous studies showed that specific altered metabolites and lipids in plasma samples varied with AD progression. (Bhawal et al. 2021) The metabolomic profile of the produced plasma patterns revealed many significantly changing metabolites in different MS types particularly RRMS and SPMS (red dots in **Supplementary Figure 13**) in all the layers. For example, it has been reported that glutamate may be related to inflammatory and neurodegenerative processes evident in MS.(Kostic et al. 2013; Pitt et al. 2000) We have also found that glucoheptonic acid metabolite is significantly different in layers one and two of the produced plasma patterns within the MagLev system. Homocysteine, another significant metabolomic marker of various neurodegenerative disorders including MS, is found in layer one of the plasma patterns.(Mititelu et al. 2006) In fact, the level of homocysteine in plasma is a good predictor of both RRMS and SPMS subtypes of MS. Although it has been reported that elevated plasma homocysteine may occur in both mild and progressive disease courses of MS, the concentration of homocysteine metabolite is significantly higher in SPMS compared with RRMS subtypes.(Oliveira et al. 2018) Presence of such metabolites (presented in **Supplementary Figure 13**) individually or in combination in particular layers of the produced plasma patterns in the MagLev system could predict various types of MS with high diagnostic value.

## 5. Conclusions

In summary, using multi-omics analysis, we have shown significant differences in the biomolecular composition of various ellipsoidal bands formed by levitating blood plasma in the MagLev system when using SPIONs as a paramagnetic medium. These biomolecular variations are, therefore, the main driver of the formation of such ellipsoidal bands. In addition, we explored the possible role of ellipsoidal bands in disease detection by comparing the biomolecular composition of each band across various subtypes of MS. Our findings reveal that among other molecular species in the plasma, lipid and metabolite profiles are a major source of variability between different MS types, while plasma proteome profiles are less discriminatory between MS patients and healthy individuals. Therefore, the MagLev technique may be used for lipid and metabolite detection for disease diagnosis through comparing the biomolecular composition of various levitated ellipsoidal bands.

## Data availability

Excel files containing the analyzed raw data are provided in SI (**Supplementary Data 1 to 7**). Mass spectrometry data will be deposited to the ProteomeXchange Consortium (http://proteomecentral.proteomexchange.org) via the PRIDE partner repository with dataset identifiers.(Vizcaíno et al. 2014)

## CRediT authorship contribution statement

**Ali Akbar Ashkarran:** Conceptualization, Methodology, Experiments, Data acquisition, Formal analysis, Project administration, Writing, reviewing & editing original draft. **Hassan Gharibi**: Conceptualization, Methodology, Formal analysis, Project administration, Writing, reviewing & editing original draft. **Dalia Abou Zeki**: Writing, reviewing & editing. **Irina Radu**: Writing, reviewing & editing. **Farnaz Khalighinejad**: Sample preparation, Writing, reviewing & editing. **Kiandokht Keyhanian**: Writing, reviewing & editing. **Christoffer K. Abrahamsson**: Writing, reviewing & editing. **Carolina Ionete**: Supervision, Project administration, Review & editing. **Amir Ata Saei**: Supervision, Project administration, Review & editing. **Morteza Mahmoudi**: Conceptualization, Supervision, Project administration, Reviewing & editing original draft, Funding acquisition.

## Acknowledgements

MM gratefully acknowledges financial support from the U.S. National Institute of Diabetes and Digestive and Kidney Diseases (grant DK131417-01). AAS was supported by the Swedish Research Council (grant 2020-00687) and the Swedish Society of Medicine (grant SLS-961262, 1086 Stiftelsen Albert Nilssons forskningsfond). We would like to thank Khashayar Afshari for his valuable discussions. Protein identification and quantification were carried out by the Proteomics Biomedicum core facility, Karolinska Institutet (https://ki.se/en/mbb/proteomics-biomedicum). Parts of the Figure 1 are adapted from Bio Render (BioRender.com).

