## Supplementary Information for "Multi-Omics Analysis of Magnetically Levitated Plasma Biomolecules"

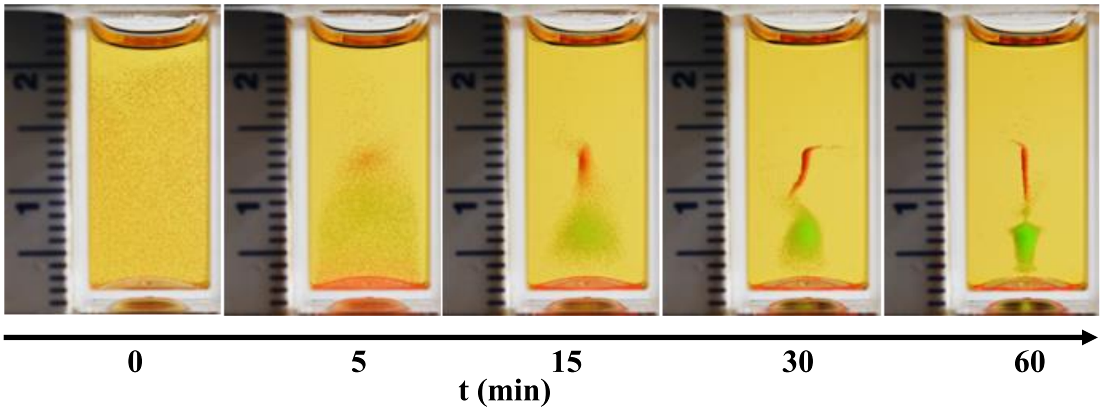


**Supplementary Figure 1.** Optical images of the levitation patterns of three mixed standard density fluorescent polyethylene microspheres (diameter of ~ 50 µm) with different densities in the MagLev system over the time.

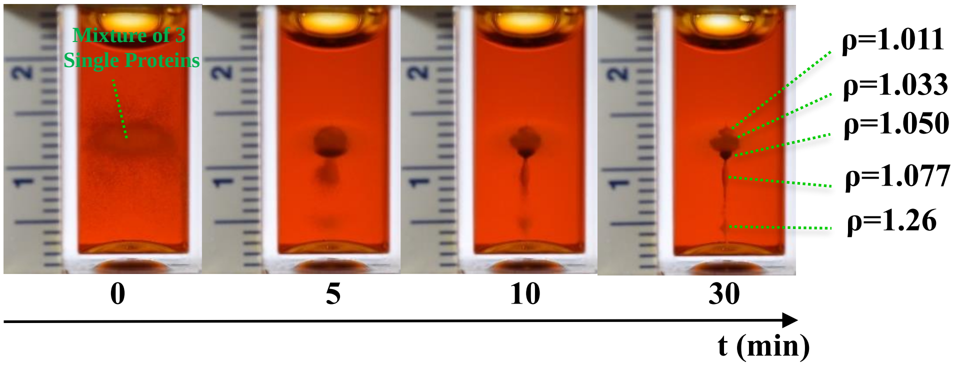


**Supplementary Figure 2.** Levitation patterns of a mixture of three mixed different molecular weights of single proteins (Lysozyme, Albumin, and Immunoglobulin G (IgG)) with five standard density fluorescent polyethylene microspheres in the MagLev system (1 mg/ml concentration of SPIONs). All density units are g/cm$^3$.

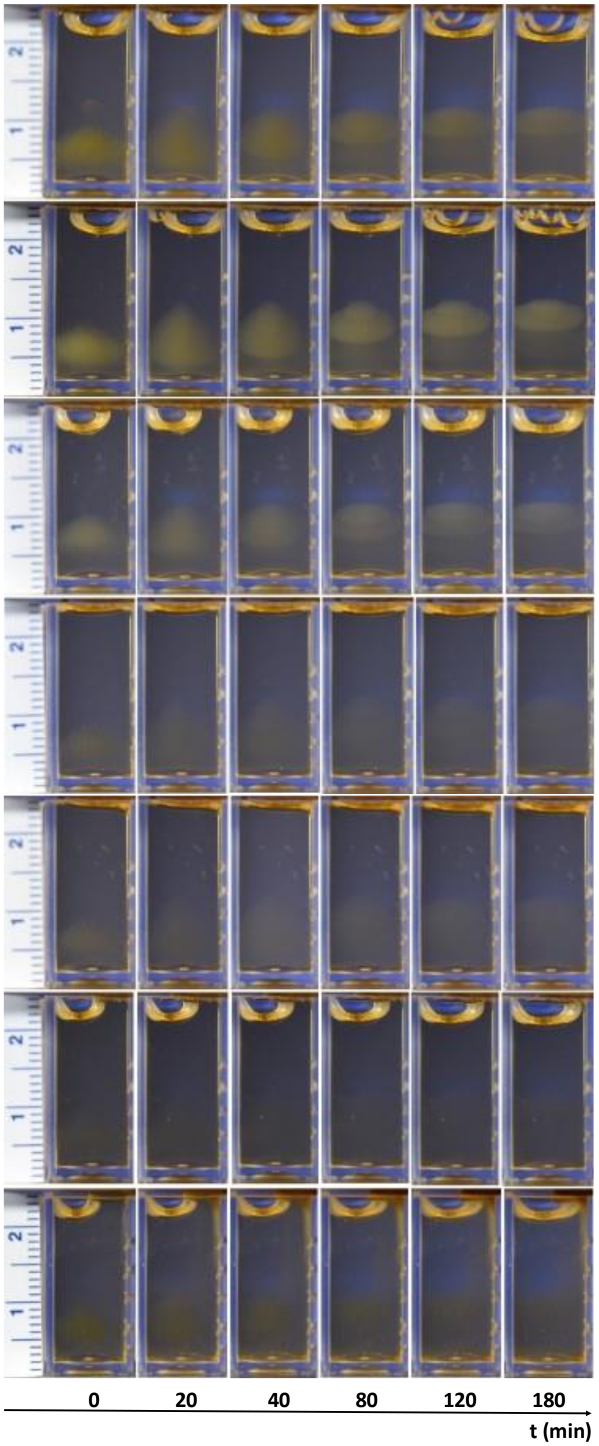


**Supplementary Figure 3.** Optical images of the levitation patterns of representative human plasma protein samples diagnosed with RRMS in MagLev at 0.06 mg/ml concentration of SPIONs.

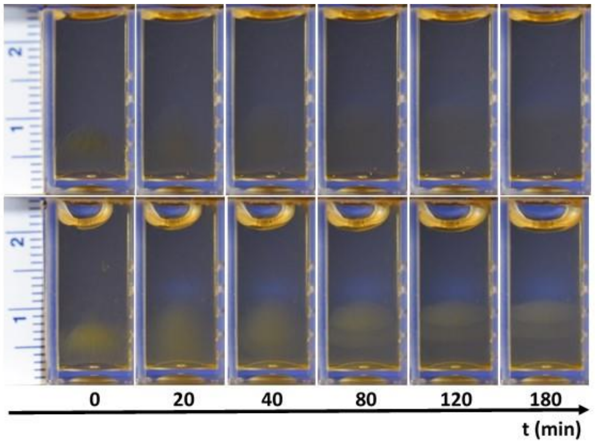


**Supplementary Figure 4.** Optical images of the Levitation patterns of representative human plasma protein samples diagnosed with PPMS in MagLev at 0.06 mg/ml concentration of SPIONs.

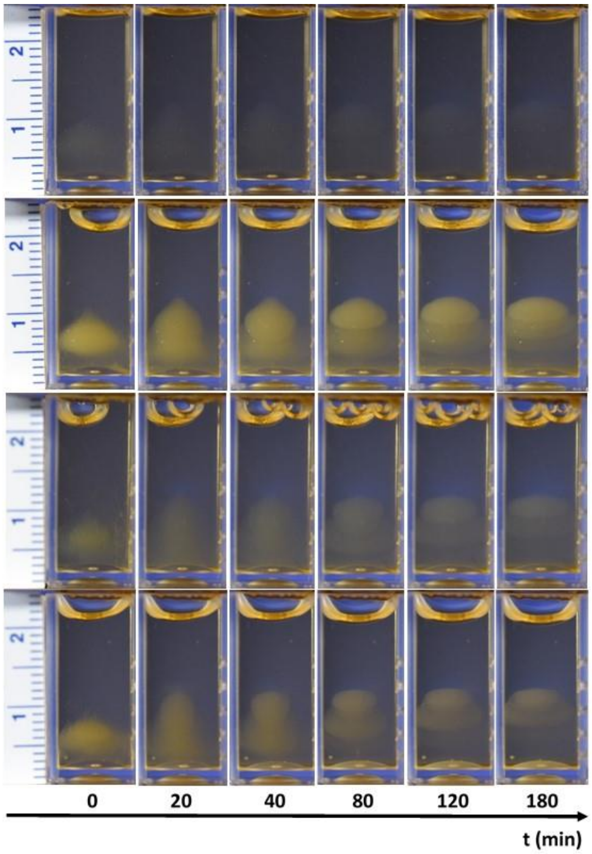


**Supplementary Figure 5.** Optical images of the Levitation patterns of representative human plasma protein samples diagnosed with SPMS in MagLev at 0.06 mg/ml concentration of SPIONs.

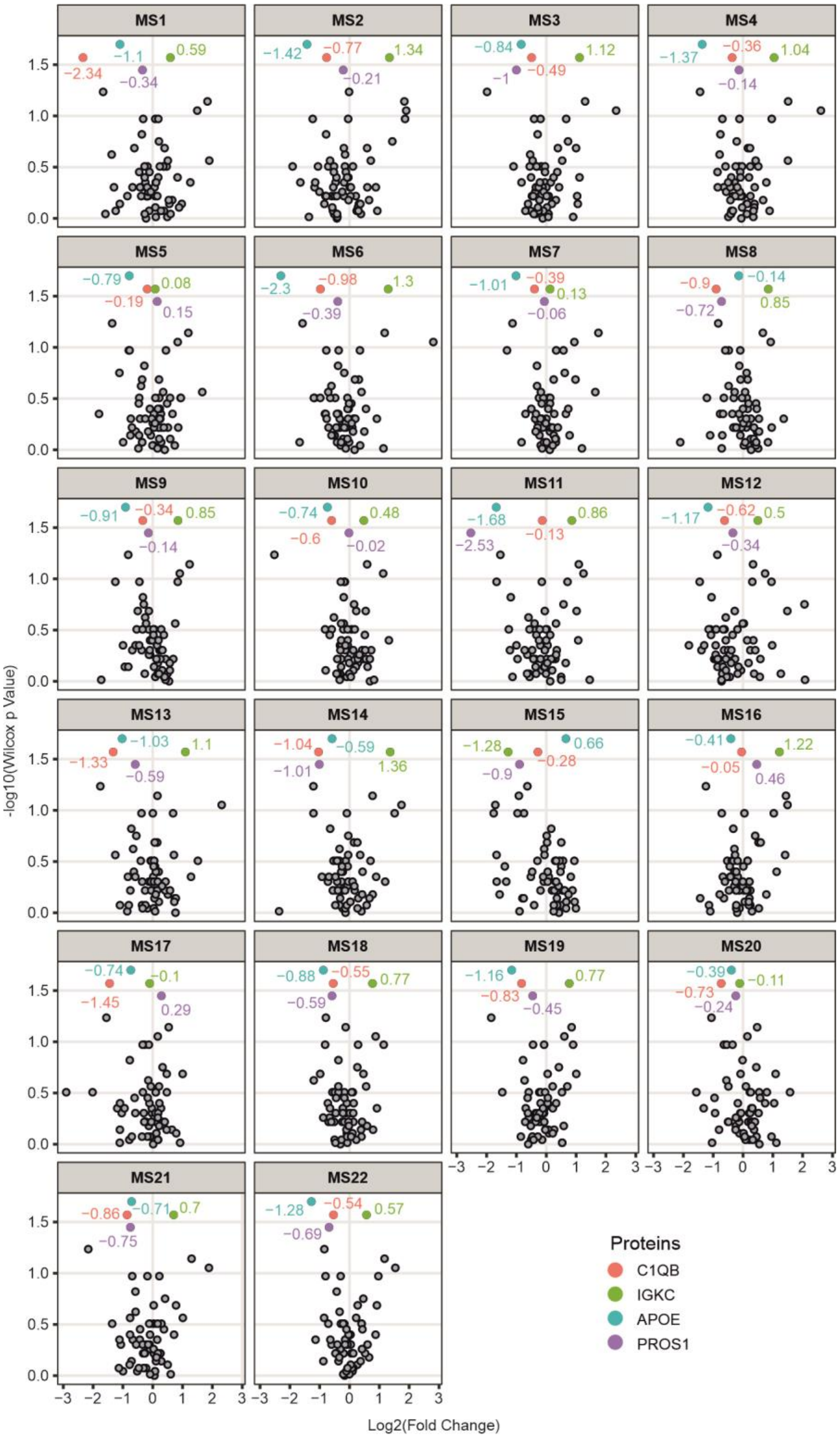


**Supplementary Figure 6. Four proteins significantly different among the healthy and MS plasma samples, shown for each MS patient vs. control.** The colored values represent log2(fold change) for each protein (MS1 to MS14 correspond to RRMS samples, MS15-MS17 correspond to PPMS samples, MS18-MS22 correspond to SPMS samples. The log2(fold changes) are shown for each protein.

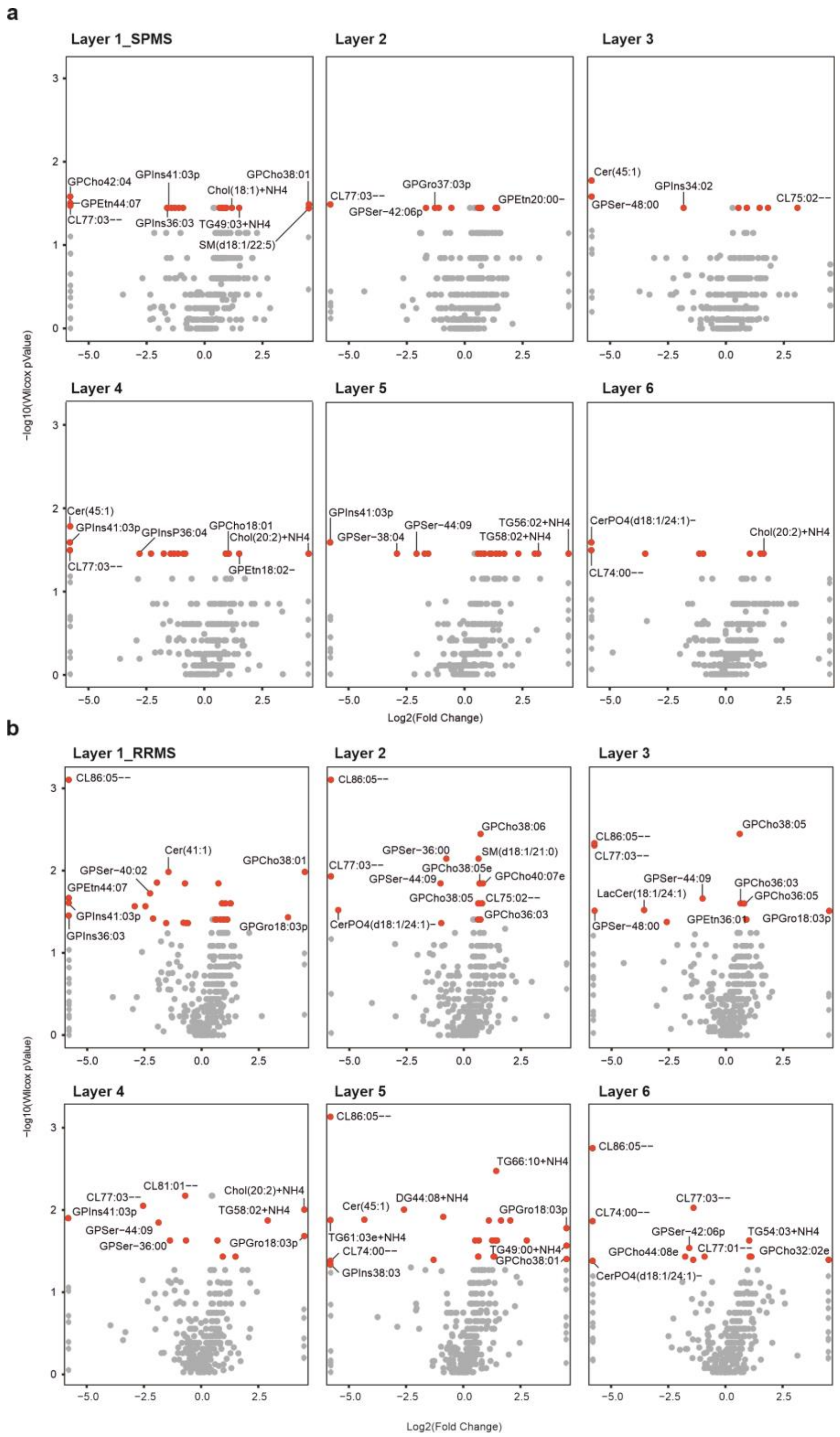


**Supplementary Figure 7. Volcano plots of lipidome profiles comparing MS samples vs. healthy in different layers.** While no significantly changing lipids were found in different layers of PPMS vs. healthy samples, there are many significantly changing lipids in the other MS types in all the layers (highlighted in red); Panel "**a**" for SPMS and panel "**b**" for RRMS types.

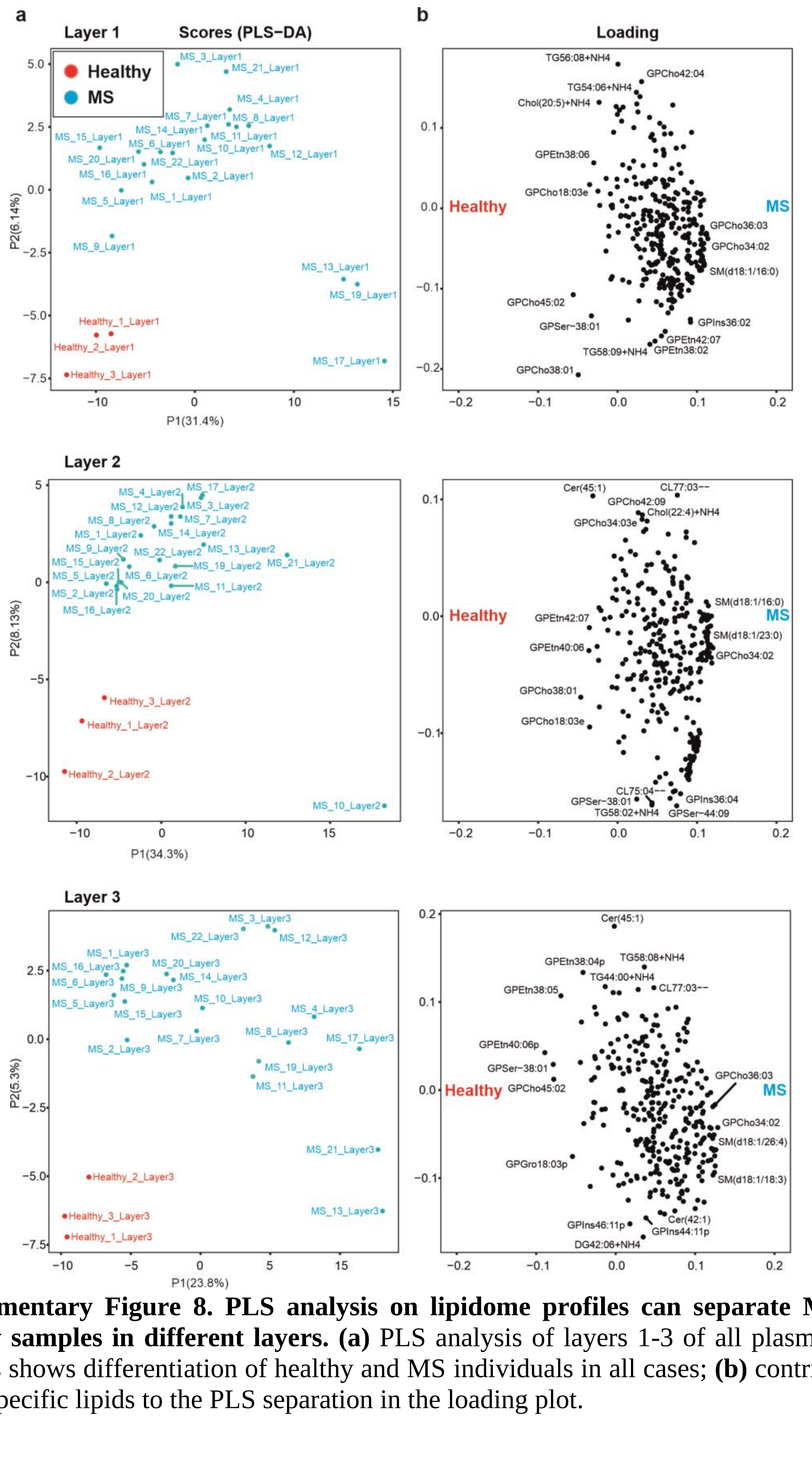

**Supplementary Figure 8. PLS analysis on lipidome profiles can separate MS vs. healthy samples in different layers. (a)** PLS analysis of layers 1-3 of all plasma lipid patterns shows differentiation of healthy and MS individuals in all cases; **(b)** contribution of the specific lipids to the PLS separation in the loading plot.

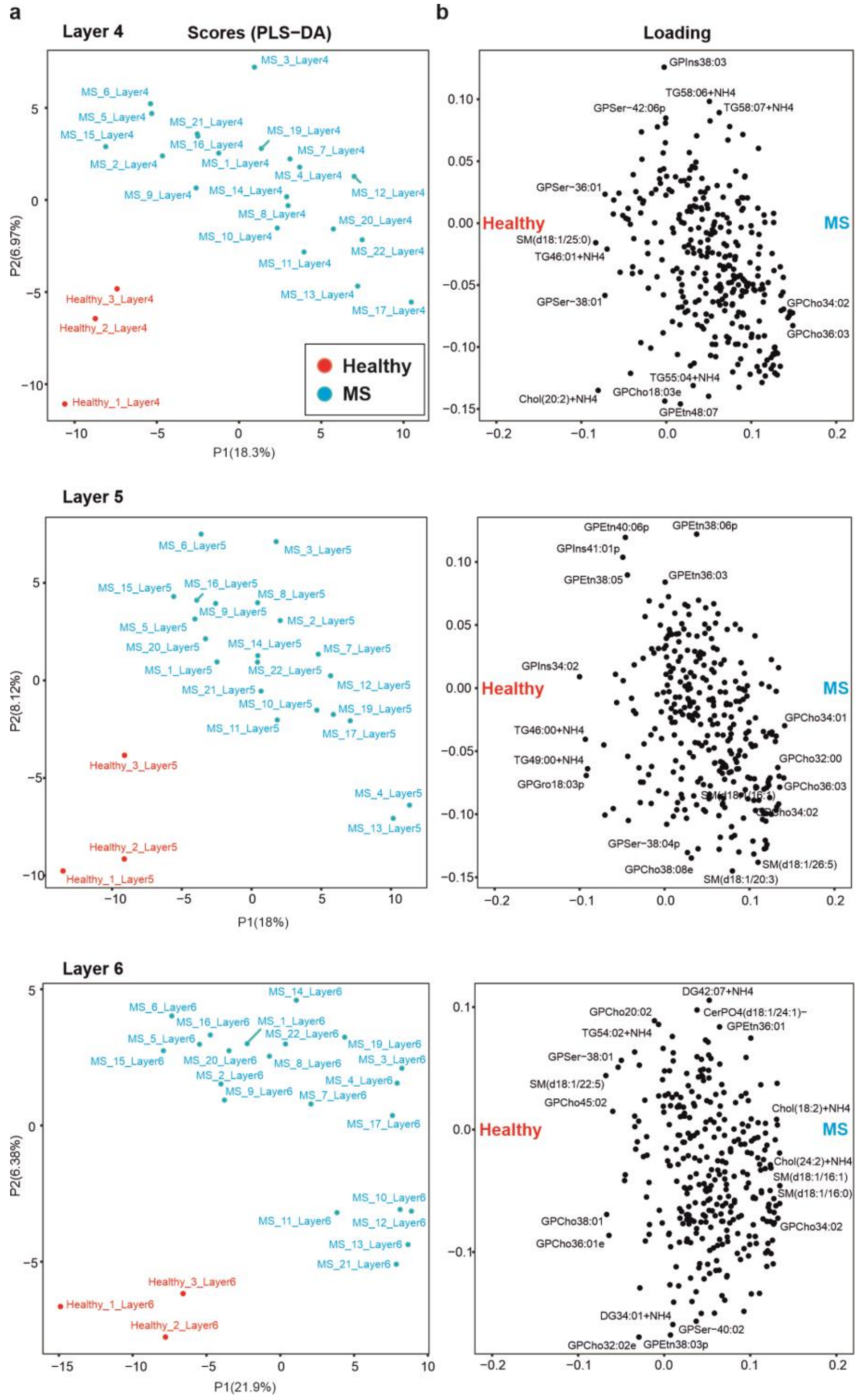


**Supplementary Figure 9. PLS analysis on lipidome profiles can separate MS vs. healthy samples in different layers. (a)** PLS analysis of layers 4-6 of all plasma lipid patterns shows differentiation of MS samples vs. healthy in all cases; **(b)** contribution of the specific lipids to the separation in the loading plot.

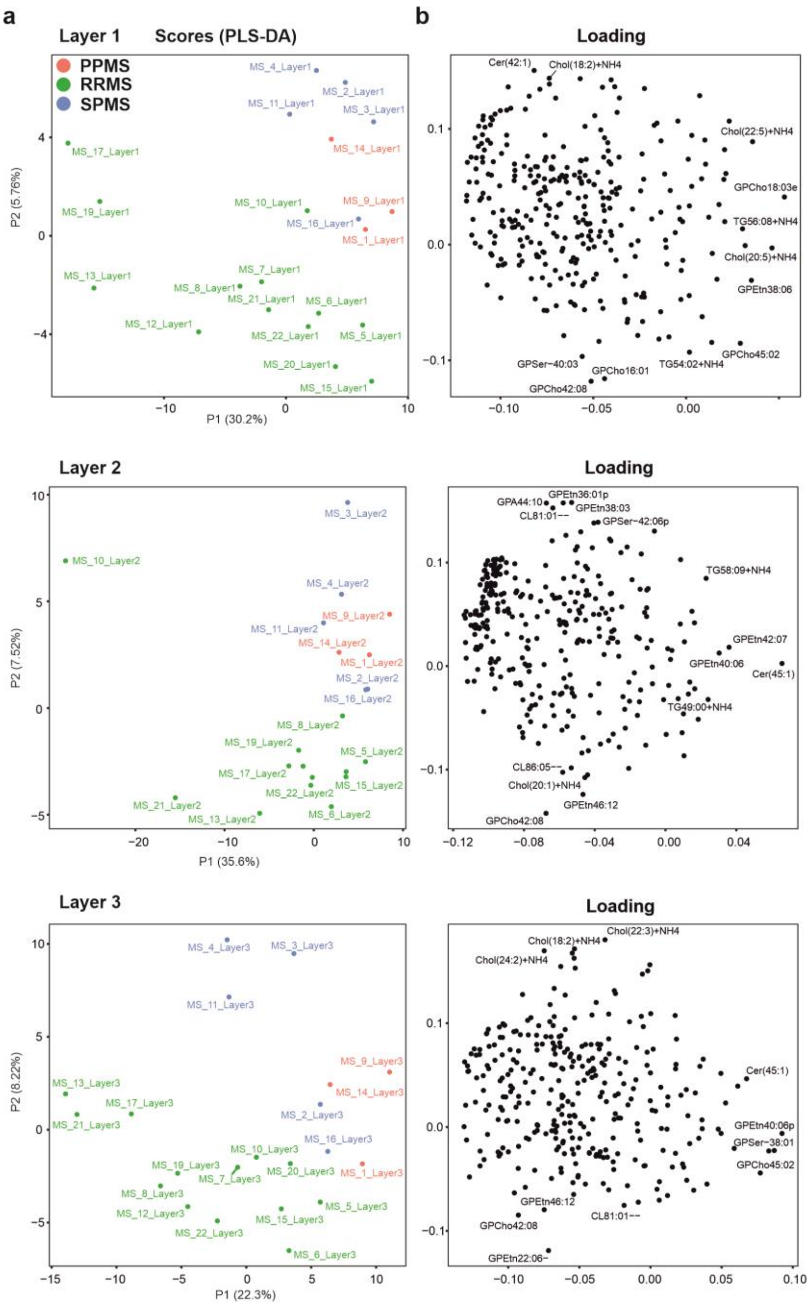


**Supplementary Figure 10. PLS analysis on lipidome profiles can separate MS types in different layers. (a)** PLS analysis of layers 1-3 of MS plasma lipid patterns shows discrimination of various types of MS; **(b)** contribution of the specific lipids to the PLS separation.

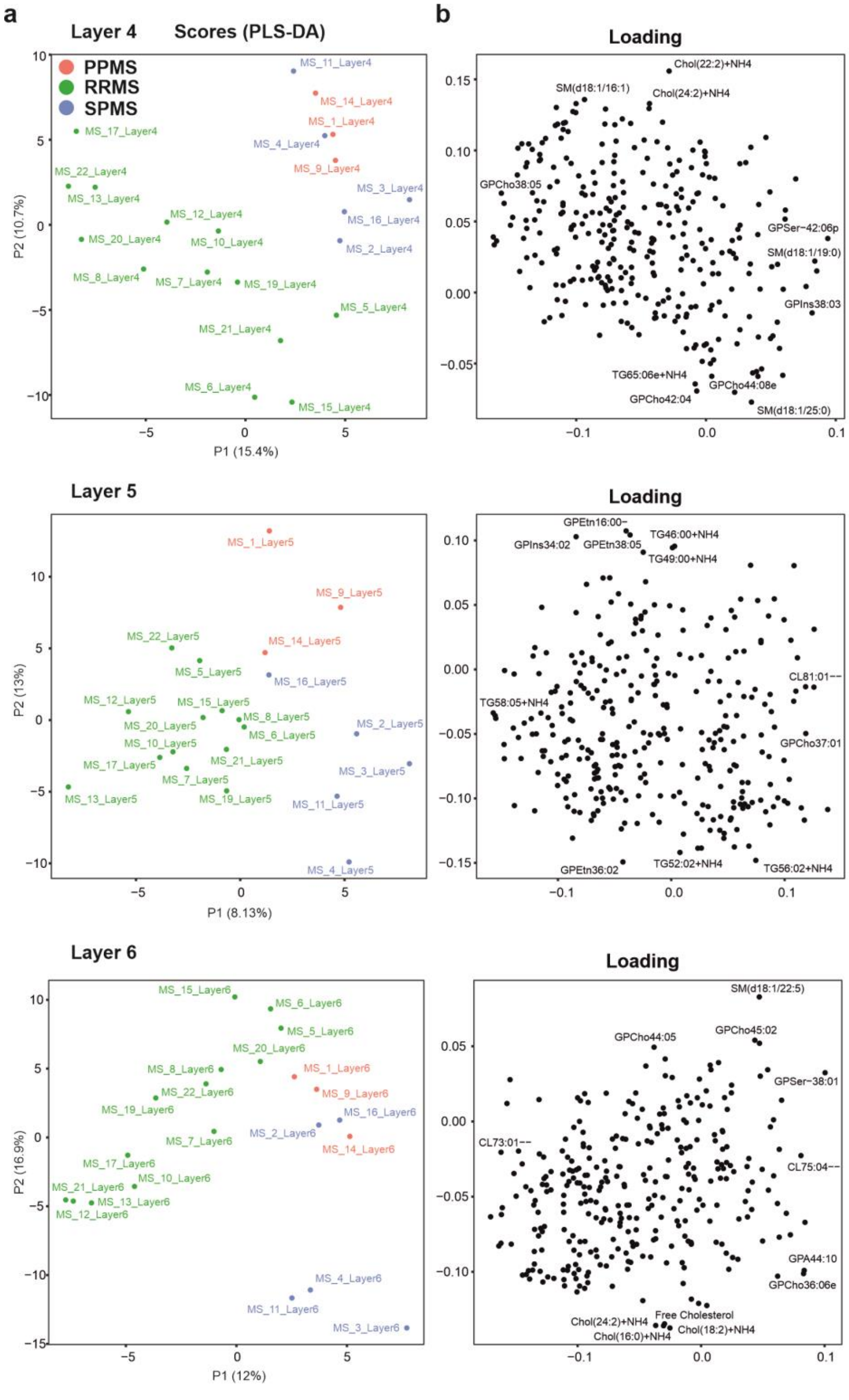


**Supplementary Figure 11. PLS analysis on lipidome profiles can separate MS types in different layers. (a)** PLS analysis of layers 4-6 of MS plasma lipid patterns shows discrimination of various types of MS; **(b)** contribution of the specific lipids to the PLS separation.

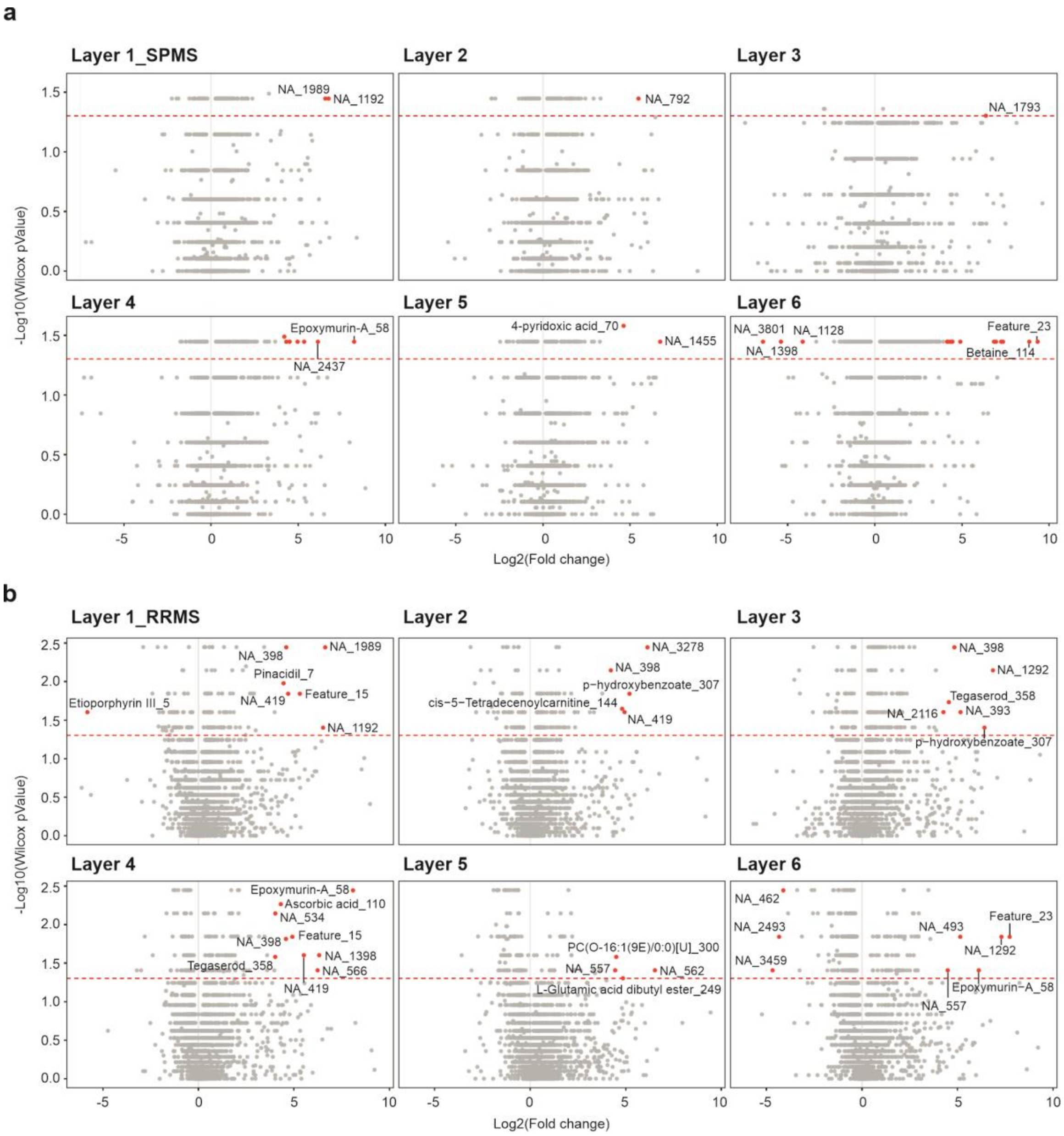


**Supplementary Figure 12. Volcano plots of metabolome profiles comparing MS samples vs. healthy in different layers.** While no significantly changing metabolites were found in different layers of PPMS MS vs. control, there are many significantly changing metabolites in the other MS types in all the layers (highlighted in red). The metabolites not identified are denoted with NA. The corresponding metabolite IDs are shown after an underscore "_" and can be found in corresponding supplementary data 7. Panel "**a**" for SPMS and panel "**b**" for RRMS subtypes.

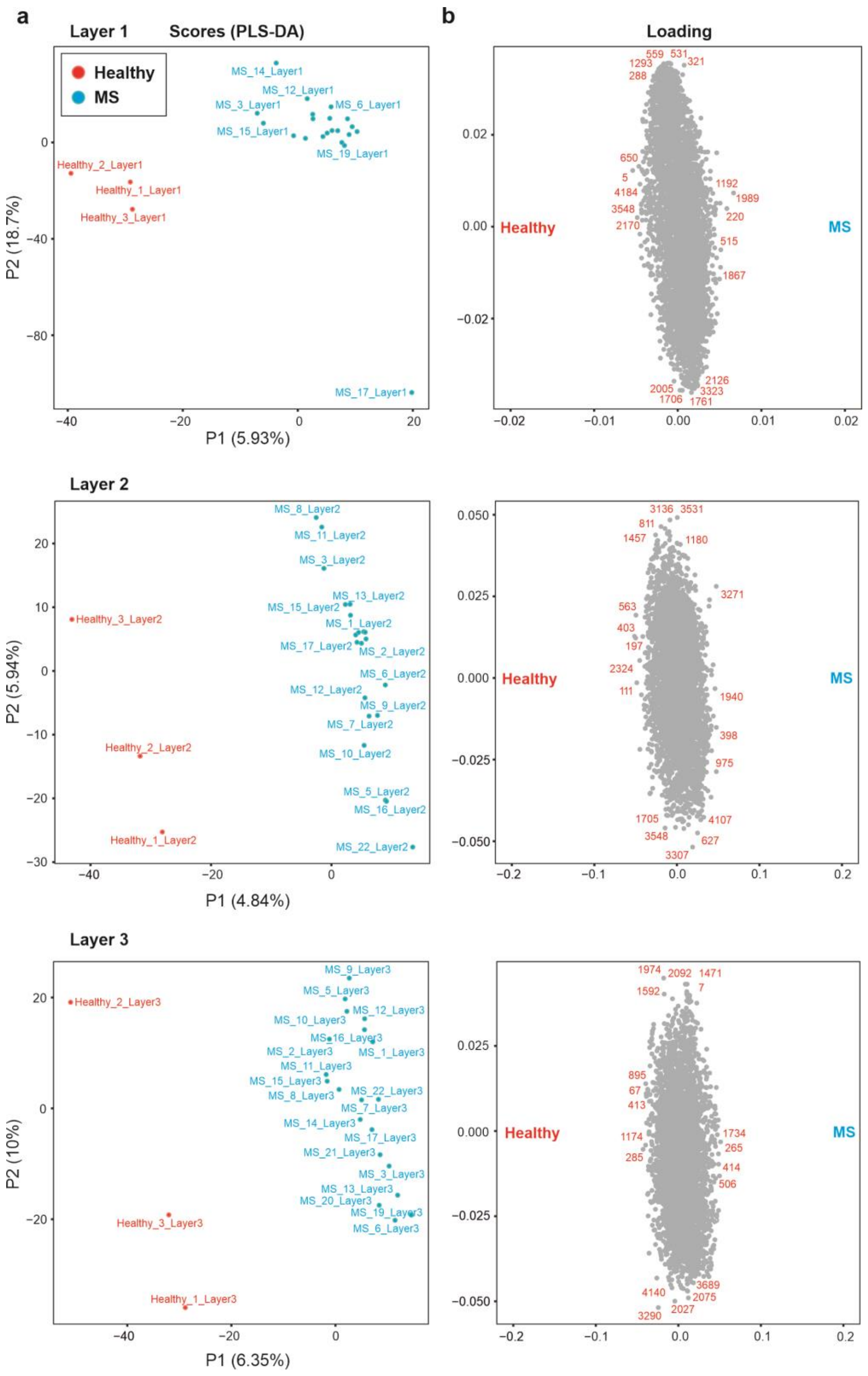


**Supplementary Figure 13. PLS analysis on metabolome profiles can separate MS vs. healthy samples in different layers.** **(a)** PLS analysis of layers 1-3 of all plasma metabolome patterns shows differentiation of healthy and MS individuals; **(b)** contribution of the specific metabolomes to the PLS separation. The corresponding metabolite IDs can be found in corresponding supplementary data 7.

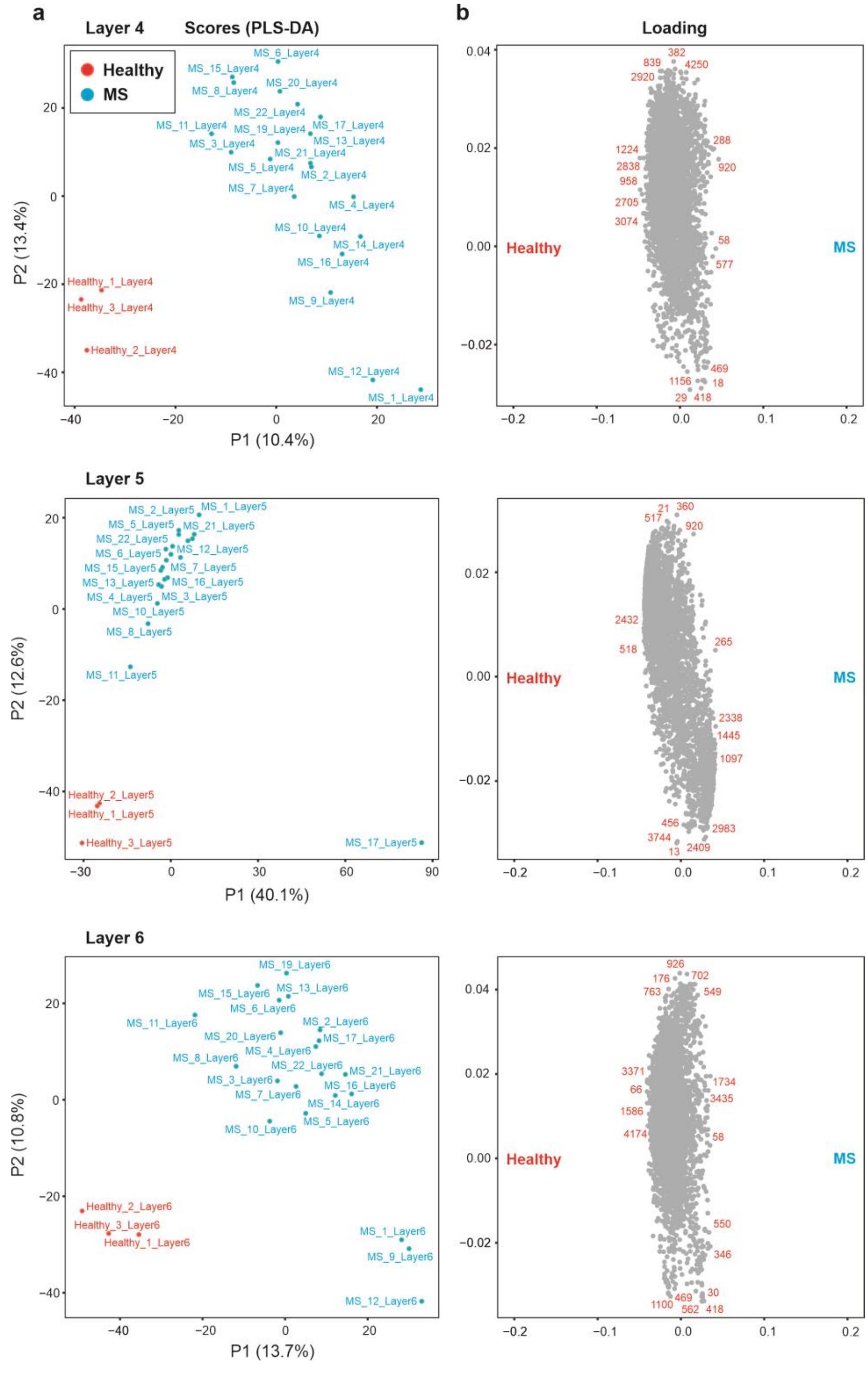


**Supplementary Figure 14. PLS analysis on metabolome profiles can separate MS vs. healthy samples in different layers. (a)** PLS analysis of layers 4-6 of all plasma metabolome patterns shows differentiation of healthy and MS individuals; **(b)** contribution of the specific metabolomes to the PLS separation. The corresponding metabolite IDs can be found in corresponding supplementary data 7.

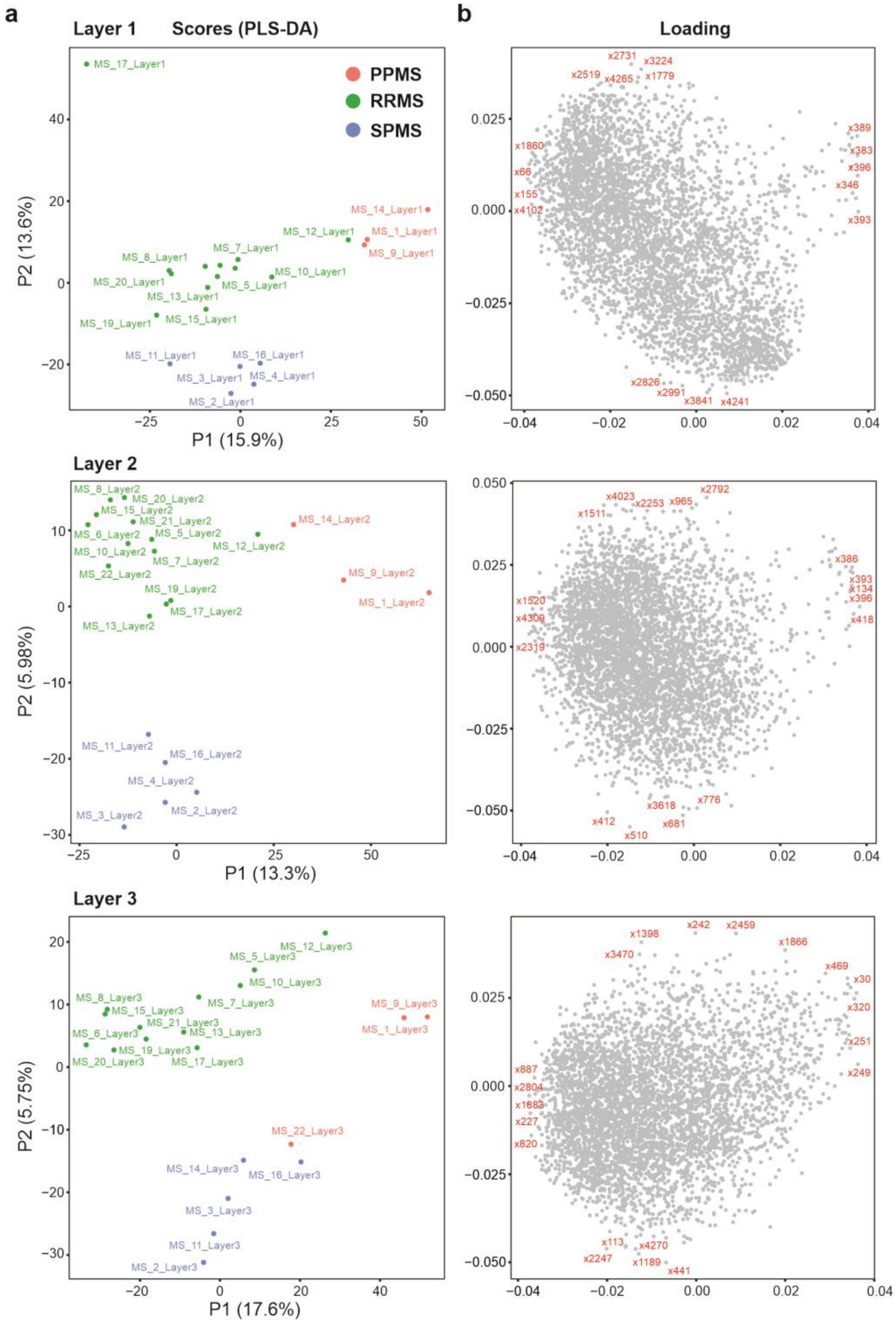

**Supplementary Figure 15. PLS analysis on metabolome profiles can separate MS types in different layers. (a)** PLS analysis of layers 1-3 of MS plasma metabolome patterns shows discrimination of various types of MS; **(b)** contribution of the specific metabolomes to the PLS separation. The corresponding metabolite IDs can be found in corresponding supplementary data 7.

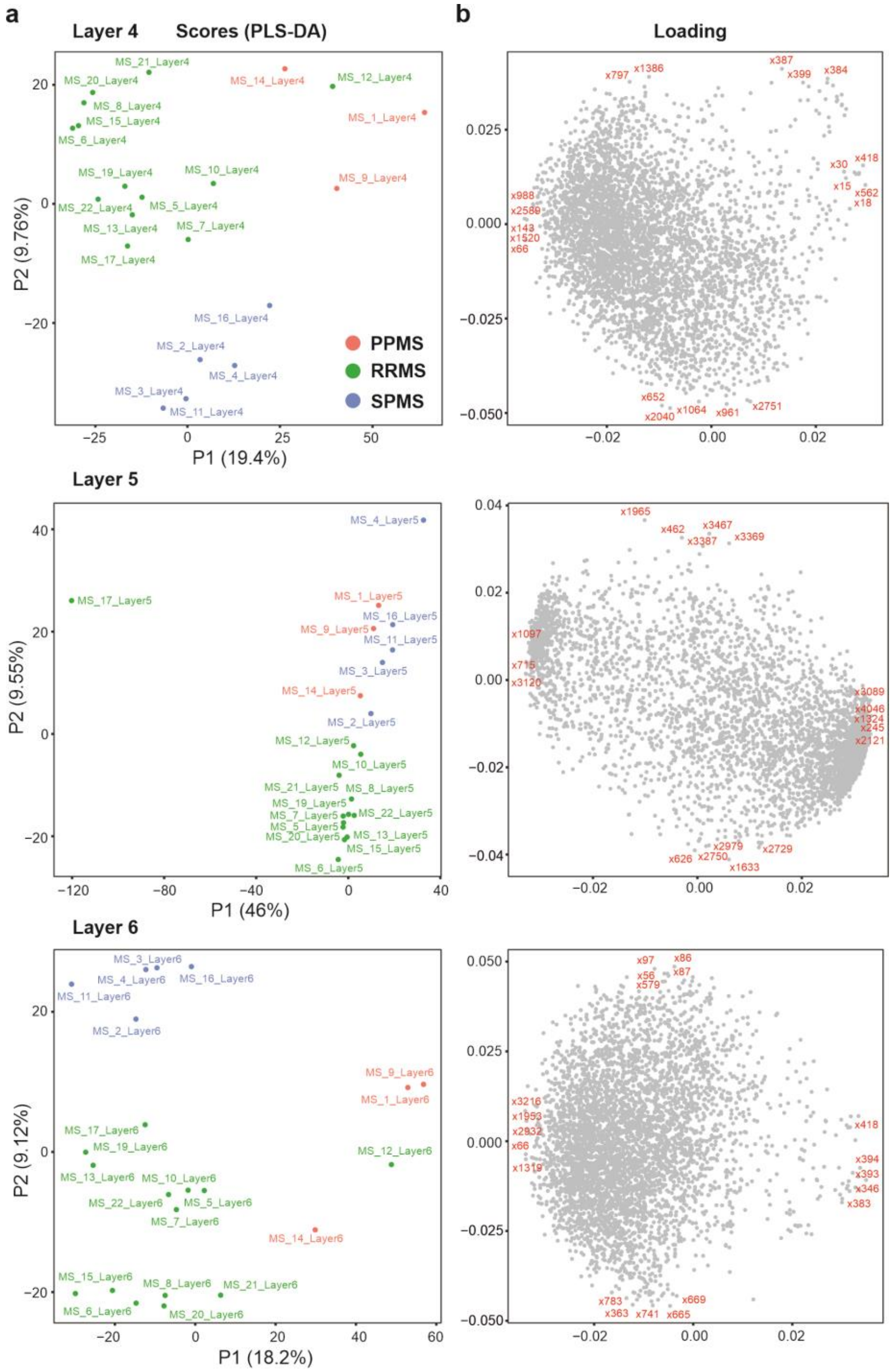


**Supplementary Figure 16. PLS analysis on metabolome profiles can separate MS types in different layers. (a)** PLS analysis of layers 4-6 of MS plasma metabolome patterns shows discrimination of various types of MS; **(b)** contribution of the specific metabolomes to the PLS separation. The corresponding metabolite IDs can be found in corresponding supplementary data 7.
